# Pathological mechanisms and candidate therapeutic approaches in the hearing loss of mice carrying human *MIR96* mutations

**DOI:** 10.1101/2024.02.20.581141

**Authors:** Morag A Lewis, Maria Lachgar-Ruiz, Francesca Di Domenico, Graham Duddy, Jing Chen, Sergio Fernandez, Matias Morin, Gareth Williams, Miguel Angel Moreno Pelayo, Karen P Steel

**Affiliations:** Wolfson Sensory, Pain and Regeneration Centre, King’s College London, Guy’s Campus, London SE1 1UL, UK; Wellcome Sanger Institute, Hinxton, CB10 1SA, UK; Servicio de Genética, Hospital Universitario Ramón y Cajal, IRYCIS and Biomedical Network Research Centre on Rare Diseases (CIBERER), km 9.100, 28034 Madrid, Spain

## Abstract

Progressive hearing loss is a common problem in the human population with no effective therapeutics currently available. However, it has a strong genetic contribution, and investigating the genes and regulatory interactions underlying hearing loss offers the possibility of identifying therapeutic candidates. Mutations in regulatory genes are particularly useful for this, and an example is the microRNA miR-96, a transcriptional regulator which controls hair cell maturation. Mice and humans carrying mutations in *Mir96* all develop hearing loss, but different mutations result in different physiological, structural and transcriptional phenotypes.

Here we present our characterisation of two lines of mice carrying different human mutations knocked-in to *Mir96*. While mice homozygous for either mutation are profoundly deaf from two weeks old, the heterozygous phenotypes differ markedly, with only one mutation resulting in hearing impairment in heterozygosis. Investigations of the structural phenotype showed that one mutation appears to lead to synaptic defects, while the other has a much more severe effect on the hair cell stereociliary bundles. Transcriptome analyses revealed a wide range of misregulated genes in both mutants which were notably dissimilar. We used the transcriptome analyses to investigate candidate therapeutics, and tested one, finding that it delayed the progression of hearing loss in heterozygous mice.

Our work adds further support for the importance of the gain of novel targets in microRNA mutants, and offers a proof of concept for the identification of pharmacological interventions to maintain hearing.

## Introduction

Progressive hearing loss is a common problem in the human population, but as yet there are no therapeutic treatments available. Part of the reason for this is that adult-onset hearing loss is a highly heterogeneous condition, and many factors can underlie an individual’s hearing impairment. However, it is known that there is a considerable genetic contribution (Cherny et al. 2020; Wingfield et al. 2007; Wolber et al. 2012), suggesting that a better understanding of the genes involved may lead to useful pathways to explore therapeutically.

Mutations affecting the microRNA miR-96 have been found to cause progressive hearing loss in humans and in mice (Lewis et al. 2009; Mencia et al. 2009; Solda et al. 2012). MicroRNAs are small noncoding RNA genes which target mRNA molecules through binding to target sites in their 3’UTR and recruiting the RISC complex to downregulate translation (reviewed in (Bartel 2004; Winter et al. 2009)). Two of the point mutations of *MIR96* that have been reported in humans cause dominant, progressive hearing loss in carriers (Mencia et al. 2009), both affecting the seed region of the microRNA, which is critical for correct binding (Lewis, Burge, and Bartel 2005). In the mouse, a third point mutation of the seed region in *Mir96^Dmdo^* results in delayed maturation of sensory hair cells in heterozygous carriers, and a complete failure of hair cells and their innervation to develop properly in homozygotes, both in the peripheral and central auditory system (Kuhn et al. 2011; Lewis et al. 2009; Schluter et al. 2018). Because of the similarity of the phenotype resulting from these three point mutations, the mechanism of action initially was thought to be the loss of the correct targets (Lewis et al. 2009; Mencia et al. 2009). However, more recent studies on mice carrying null alleles of either *Mir96* and the nearby *Mir183* (Lewis et al. 2020) or of all three microRNAs of the *Mir183* family (*Mir96*, *Mir183* and *Mir182*) (Fan et al. 2017; Geng et al. 2018) found that heterozygous carriers of the null alleles have no hearing phenotype, suggesting that the gain of novel targets due to the changed seed sequence resulting from point mutations is important in the *Mir96* mutant phenotype, not just the loss of normal targeting. Transcriptome analyses of the *Mir96^Dmdo^* mutant organ of Corti showed that miR-96 controls a broad regulatory network (Lewis et al. 2016), suggesting that a better understanding of the core genes – particularly the direct targets of miR-96 – may suggest candidate therapeutic targets.

Here we present data from mice carrying the two seed region point mutations reported in human families (Mencia et al. 2009). Both mutations result in profound deafness in homozygotes, but the heterozygote phenotype differs, further supporting the importance of the gain of novel targets in microRNA mutant phenotypes. We have investigated the structural and transcriptomic changes, and found evidence for the involvement of different pathological mechanisms; one mutation appears to affect the synapses while the other has a more severe effect on the hair bundle. Finally, we have demonstrated the potential for use of these data to identify candidate therapeutics, and we have tested and confirmed the effect of one such candidate to delay the progression of hearing loss seen in heterozygotes.

## Materials and methods

### Ethics statement

Mouse studies were performed in compliance with UK Home Office regulations and the Animals (Scientific Procedures) Act of 1986 (ASPA) under UK Home Office licencing, and the study was approved by the King’s College London Ethical Review Committee. Mice were culled using methods permitted under these licences to minimise any possibility of suffering.

### Mouse generation and maintenance

*Mir96* mutant mice were generated by the Mouse Genetics Project at the Wellcome Sanger Institute by inserting targeted mutations in mouse ES cells in a C57BL/6N genetic background, after which the selection cassette was removed using Flp recombinase. The mouse lines used are *Mir96^tm2.1Wtsi^* (hereafter referred to as *Mir96^+13G>A^*, corresponding to human family s403 (Mencia et al. 2009; Modamio-Hoybjor et al. 2004), ES cell line BEPD0019_D04) and *Mir96^tm3.1Wtsi^* (hereafter referred to as *Mir96^+14C>A^*, corresponding to human family s1334 (Mencia et al. 2009), ES cell line BEPD0003_D07) (Fig 1). Both mouse lines will be available through the European Mouse Mutant Archive (EMMA). Both colonies were generated and maintained on a C57BL/6N genetic background and were maintained by heterozygous intercrosses or homozygotes crossed with heterozygotes, and both mutant lines produce viable and fertile homozygous offspring. For the testing of the *Mir96^+13G>A^* allele on the C3HeB/FeJ background, *Mir96^+13G>A^* homozygotes were crossed to C3HeB/FeJ mice, producing F1 mice which were heterozygous on a 50% C3HeB/FeJ, 50% C57BL/6N background. These F1 mice were then bred together, and ABR measurements were taken from their F2 offspring (wildtype, heterozygous and homozygous for the *Mir96^+13G>A^*allele, on a 50% C3HeB/FeJ, 50% C57BL/6N background).

**Figure 1.**
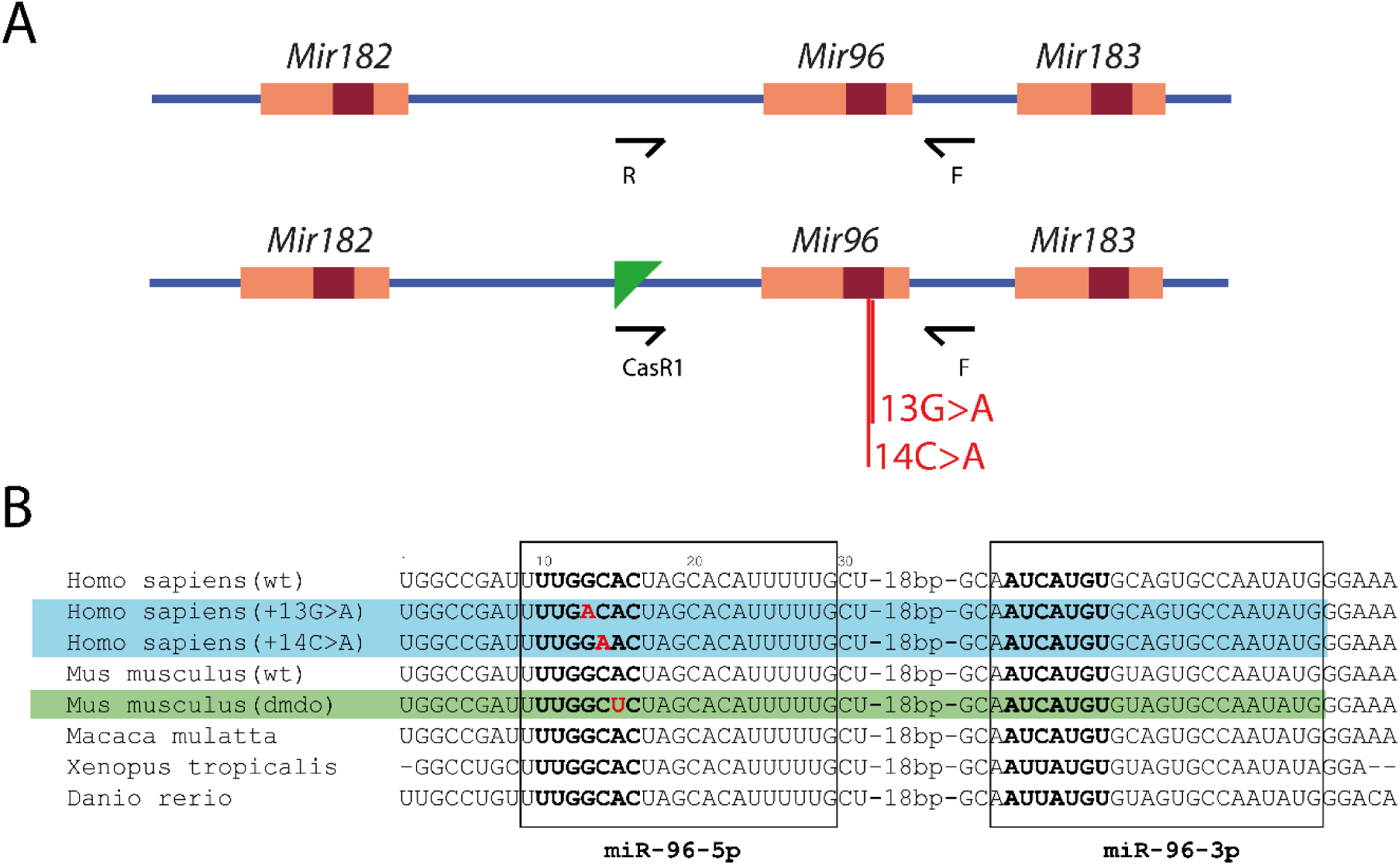
Schematic showing the allele structure of the mice used in this work. A) Allele structure and primer location for the wildtype (top) and mutant (bottom) alleles. *Mir96* is one of three microRNAs in the *Mir-183/96/182* gene cluster. Maroon boxes indicate the seed regions, and the position of the mutations are labelled. Arrows indicate the location and orientation of the primers used for genotyping. Green triangle: FRT site. Not to scale. B) Partial sequence of the *Mir96* stem-loop showing the point mutations associated with deafness in mice and humans; the sequence shown is from the start to the end of the human gene sequence but not all the species share the same start and end points. The boxes indicate the mature microRNA sequences. The seed region, critical for the correct identification of target mRNAs, is shown in bold. Mutations are shown in red. The two human mutations carried by the *Mir96^+13G>A^* and *Mir96^+14C>A^*mutants are highlighted in blue; the numbers refer to the position from the start of the human gene. The point mutation in the *Mir96^Dmdo^* mouse is highlighted in green (Lewis et al. 2009; Mencia et al. 2009).

### Genotyping

*Mir96^+13G>A^* and *Mir96^+14C>A^* mice were genotyped by PCR using template DNA extracted from tissue. The mutant allele was amplified using a gene-specific forward primer (5’- CTCACCCCTTTCTGCCTAGA-3’) paired with a reverse primer designed against the remaining FRT site left after removal of the selection cassette (5’- TCGTGGTATCGTTATGCGCC-3’). The wildtype allele was genotyped in a similar way using the same forward primer combined with a reverse primer specific for the wildtype allele (5’- CCTAAGGTAAGCCACTGATGG -3’). The resulting band sizes were 588 bp for the wildtype product and 587 bp for the mutant product. The location of the primers with respect to *Mir96* and the cassette is shown in Fig 1.

### Auditory Brainstem Response

The hearing of wildtype and mutant *Mir96^+13G>A^* and *Mir96^+14C>A^* mice was tested using the Auditory Brainstem Response (ABR) as described in (Ingham, Pearson, and Steel 2011). Briefly, mice were sedated using a ketamine/xylazine mix (10mg ketamine and 0.1mg xylazine in 0.1ml per 10g body weight), and responses were recorded from three subcutaneous needle electrodes. The reference electrode was placed over the left bulla, the ground over the right bulla, and the active electrode was placed on the top of the head. Responses were recorded from 256 stimulus presentations per frequency (broadband click and 3kHz, 6kHz, 12kHz, 18kHz, 24kHz, 30kHz, 36kHz and 42kHz pure tone frequencies at sound levels from 0-95dB, in 5dB steps). Mice were then recovered using atipamezole (0.01mg atipamezole in 0.1ml per 10g body weight). The threshold for each frequency is the lowest intensity at which a waveform could be distinguished, and this was identified using a stack of response waveforms.

### Noise exposure

8 week-old heterozygous and wildtype *Mir96^+13G>A^* mice were subjected to 8-16kHz octave-band noise at 100dB SPL for 1 hour while awake and unrestrained in separate compartments within an exposure chamber set up to provide a uniform sound field (Holme and Steel 2004), as described in (Ingham et al. 2020; Lewis et al. 2020). ABRs were carried out 1 day prior to noise exposure, and 1 day, 3 days, 7 days, 14 days and 28 days after exposure. Sham controls were littermates who spent the same time in the exposure chamber without the noise, and went through the same set of ABR measurements at the same time.

### Scanning electron microscopy

The cochleae of wildtype, heterozygous and homozygous mice at P28 were collected and fixed for two hours in 2.5% glutaraldehyde in 0.1M sodium cacodylate buffer with 2mM CaCl_2_. Samples were then fine dissected in PBS to expose the organ of Corti and processed according to the osmium tetroxide-thiocarbohydrazide (OTOTO) method (Hunter-Duvar 1978) before dehydration through an ethanol series, critical point drying, and mounting. Samples were gold-coated to 4nm thickness. Regions of the cochlea were identified using the frequency-place map described by (Muller et al. 2005). Images were taken using a JEOL JSM 7800 Prime scanning electron microscope. A standard magnification of 60x was used to view the whole length of the organ of Corti, and higher magnifications were used for close-ups on hair cell rows (2000x) and individual hair cells (15000-23000x).

### Whole-mount dissection and immunohistochemistry

The cochleae of 4 week-old mice were fixed in 4% paraformaldehyde (PFA) for 2 h at room temperature (RT). After washing in PBS, samples were decalcified by overnight incubation with 0.1M EDTA disodium salt (Reagecon, cat.no. ED2015) at RT. Following dissection of the organ of Corti, samples were permeabilised with 5% Tween in PBS for 30 min and blocked with 0.5% Triton X-100 and 10% Normal Horse Serum (NHS) in PBS for 2h at RT. The samples were incubated with the primary antibodies diluted in a 1:2 blocking solution in PBS overnight at 4°C.

*Mir96^+13G>A^* and *Mir96^+14C>A^*samples were stained for the presynaptic marker C-terminal-binding protein 2 (CtBP2), the postsynaptic marker glutamate receptor 2 (GluR2) and the hair cell marker Myo7a. The primary antibodies used were rabbit anti-Myosin VIIa (1:200, 25-6790, Proteus), mouse IgG2 anti-GluR2 (diluted 1:2000, MAB397, Emd Millipore) and mouse IgG1 anti-CtBP2 (diluted 1:200, 612044, BD Transduction Laboratories). The day after, samples were washed three times with PBS and incubated for 1h with the secondary antibodies Alexa Fluor 647-conjugated chicken anti-rabbit (1:200, #A21443, Life Technologies), Alexa Fluor 488-conjugated goat anti-mouse (IgG2a) (diluted 1:1000, #A21131, Life Technologies) and Alexa Fluor 568-conjugated goat anti-mouse (IgG1) (1:1000, #A21124, Life Technologies). This step was followed by a second incubation under the same conditions with fresh secondary antibody. Finally, specimens were rinsed with PBS and mounted using ProLong Gold Antifade Mountant with DAPI (P36931, Life Technologies) and stored at 4°C.

### Confocal imaging, synapses, and hair cell quantification

Samples were imaged with a Zeiss Imager 710 confocal microscope interfaced with ZEN 2010 software. All images were captured with the plan-Apochromat 63x/1.4 and 40x/1.4 Oil DIC objectives. Overview images of all samples were captured using a lower magnification (x10) to image all pieces of the whole organ of Corti for frequency-place mapping. The Fiji Measure_line plugin (Eaton Peabody Laboratories, https://www.masseyeandear.org/research/otolaryngology/eaton-peabody-laboratories/histology-core) was used to map cochlear length to cochlear frequencies. This plugin is based on the mouse tonotopic cochlear map described by Müller and colleagues (Muller et al. 2005).

To image the synaptic puncta in *Mir96^+13G>A^* and *Mir96^+14C>A^* samples, two non-overlapping images were acquired at the 12 kHz best-frequency region. A 2.0 optical zoom was used and a z-step of 0.25 µm was used to ensure that all synaptic puncta were imaged. Images containing puncta were merged in a z-stack and the z-axis projection was used to quantify synapses, which were defined as the colocalization of CtBP2 and GluR2-labelled puncta. Synapse quantification was performed manually using the cell counter plugin in Fiji. The total number of ribbon synapses was divided by the number of IHCs present in the image (Myo7a-labelled) to determine the number of ribbon synapses per IHC. In the cases where an IHC was only partially visible in the image, the synapses corresponding to that cell were not counted.

### Retrotranscription and quantitative PCR (RT-qPCR)

The organs of Corti of postnatal day (P)4 mice were dissected and stored at -20°C in RNAprotect Tissue Reagent (RNAlater®) (QIAGEN, cat. no. 76106). Total RNA was extracted using the SPLIT RNA extraction kit (Lexogen) according to the manufacturer’s instructions, quantified in a Nanodrop spectrophotometer and normalised to the same concentration within each litter. SuperScript™ VILO™ cDNA Synthesis Kit (Thermo Fisher Scientific, cat. no. 11754050) was used for DNase treatment and cDNA synthesis according to the manufacturer’s instructions.

Quantitative RT-PCR was performed in a CFX Connect Real-Time PCR Detection System (Bio-Rad Laboratories, Cat. No. 1855201) using TaqMan probes [Applied Biosystems: *Hprt* (Mm01545399_m1), *Prox1* (Mm00435971_m1), *Ocm* (Mm00712881_m1) and *Slc26a5* (Mm01167265_m1)] and SsoAdvanced Universal SYBR Green Supermix (Bio-Rad, #1725284). Three technical replicates of each sample and probe were carried out. The number of biological replicates (mice) tested per probe is indicated in the figure legends.

Relative expression levels were calculated using the 2^−ΔΔCΤ^ method (Livak and Schmittgen 2001). The calibrator was the wildtype littermate C_t_ for the same probe, with *Prox1* as an internal control for sensory tissue, because it is expressed in supporting cells (Bermingham-McDonogh et al. 2006).

### RNA-seq and data analysis

Cochleae were dissected and RNA was extracted as described above. Six sex-matched wildtype and homozygous mutant littermate pairs were analysed. Samples were collected within the same 1.5 h time window (from 6 hours after lights on) to control for the effects of circadian rhythms on gene expression. The age selected for RNA-seq was postnatal day 4 (P4), as it had been previously described that hair cells are still present at P4 in *Mir96^Dmdo^* mice, and to allow comparison with previous transcriptomic data from *Mir96^Dmdo^* (Lewis et al. 2020; Lewis et al. 2009). The RNA was extracted using the SPLIT RNA extraction kit (Lexogen) according to the manufacturer’s instructions. RNA was quantified using the Qubit RNA HS Assay Kit (Thermo Fisher Scientific Cat.#Q32855) and its quality was assessed on Agilent RNA 6000 Nano or Pico Chips (Agilent Technologies, Cat.# 5067-1511 or Cat.# 5067-1513) before proceeding to library preparation.

Library generation and sequencing were performed at the Center for Cooperative Research in Biosciences (CIC bioGUNE; Madrid) following the “TruSeq Stranded mRNA Sample Preparation Guide” with the corresponding kit [Illumina Inc. Cat.# RS-122-2101 or RS-122-2102] and sequenced on an Illumina HiSeq 4000 machine as paired-end 101bp reads. RNA-seq data was analysed using an in-house designed pipeline, as follows. Raw data were pre-processed to mask undetermined nucleotides based on quality using the FastX Toolkit (http://hannonlab.cshl.edu/fastx_toolkit/index.html) and Trimmomatic 0.39 for adapter trimming based on the sequence. Hisat 2.1.0 (Kim, Langmead, and Salzberg 2015) was used to align the reads against the GRCm39 reference genome. The Samtools package was used for SAM to BAM conversion (Danecek et al. 2021) and the count matrixes were generated by htseq v0.11.2 (Anders, Pyl, and Huber 2015). Finally, the edgeR package (Robinson, McCarthy, and Smyth 2010) was used to perform a generalised linear model likelihood ratio test. The resulting lists of genes differentially expressed between homozygotes and wildtype samples are shown in Tables S1 and S2.

To study the direct effect of the mutant miR-96 on the transcriptomes of the homozygous mutants, we used Sylamer (van Dongen, Abreu-Goodger, and Enright 2008), which plots over- and underrepresentation of nucleotide words of specific length within the 3’UTRs of a gene list, in our case the lists presented in Tables S1 and S2. To study the indirect effects, we used Gene Set Enrichment Analysis (GSEA), to compare the transcriptomes to previously-defined gene sets and identify any enrichment in either genotype (Mootha et al. 2003; Subramanian et al. 2005). The GSEA Preranked option (GSEA v4.3.2) was used to run the gene set enrichment analysis against the lists of differentially expressed genes ranked by log fold change (logFC) (ordered from the most upregulated to the most downregulated genes in the homozygote samples), and to determine whether any gene sets were enriched at either end of our ranked gene list. The gene sets used in this study were: hallmark gene sets, canonical pathways gene sets derived from the Reactome pathway database, regulatory targets gene sets (potential targets of regulation by transcription factors or microRNAs) and gene ontology gene sets (biological processes, BP; cellular components, CC; and molecular function, MF). All the gene sets were downloaded from the Molecular Signature Database (MSigDB v7.1, https://www.gsea-msigdb.org/gsea/msigdb/index.jsp), a collection of annotated gene sets for use with GSEA software.

The GSEA output was uploaded into Cytoscape for visualization and interpretation of enrichment analysis results using the tools EnrichmentMap for network visualization and WordCount and AutoAnnotate for interpretation and to define the major biological clusters, as described in (Reimand et al. 2019).

### Amitriptyline delivery

200µg/ml amitriptyline hydrochloride (Abcam, cat.no. ab141902) was provided in drinking water with 2% w/v saccharin sodium salt hydrate (Scientific Laboratory Supplies, cat.no. s1002-1kg) to increase palatibility, as described in (Caldarone et al. 2003). Mice were first moved to saccharin in drinking water, then after acclimatisation, they were moved to amitriptyline. Control mice were maintained on the saccharin-only water. Their bodyweight was monitored daily to ensure they remained healthy during acclimatisation. They were then set up in breeding pairs. Offspring were maintained on the same water as their parents for ABR testing up to 6 months old.

### Statistical analyses

A one-way ANOVA with Tukey’s correction for multiple comparisons was used to compare the three different experimental groups (homozygotes, heterozygotes, and wildtypes) in the *Mir96^+13G>A^* and *Mir96^+14C>A^* synapse quantification, and to generate multiplicity-adjusted p-values. 10 to 14 hair cells were counted in each frequency region.

For the qPCR, wildtype and homozygote groups were compared using the Wilcoxon rank sum test, because it is a suitable test for small sample sizes and populations of unknown characteristics (Bridge and Sawilowsky 1999).

ABR data were first transformed using the arcsine transformation, and then analysed as described in (Lewis et al. 2020). This analysis uses separate linear models for each frequency with a compound symmetric covariance structure and restricted Maximum Likelihood Estimation (Duricki, Soleman, and Moon 2016), and permits the inclusion of all data, unlike the repeated measures ANOVA, which cannot include partial data (for example, if a mouse dies before completion of the full set of ABR measurements) (Krueger and Tian 2004). For each stimulus the interaction of all variables was measured (genotype and age, and noise and drug administration where relevant), followed by Bonferroni correction for multiple testing, using SPSS v25 (IBM).

FDR-corrected p-values were calculated for the RNAseq data by applying the Benjamini-Hochberg method to the p-values generated by edgeR (Robinson, McCarthy, and Smyth 2010).

## Results

### *Mir96^+13G>A^* heterozygous mice have normal hearing while *Mir96^+14C>A^* heterozygous mice exhibit progressive hearing loss

Auditory Brainstem Response (ABR) recordings showed that *Mir96^+13G>A^*and *Mir96^+14C>A^* homozygotes exhibit profound deafness at all ages tested, showing no response at the highest sound level tested (95 dB sound pressure level (SPL)) at any of the ages studied (14 days to 6 months old) (Fig 2). *Mir96^+14C>A^* heterozygous mice have mild progressive hearing loss most pronounced at high frequencies and progressing with age to lower frequencies (Fig 2), which correlates with the human phenotype (Mencia et al. 2009; Modamio-Hoybjor et al. 2004). However, *Mir96^+13G>A^* heterozygous mice have normal hearing up to 6 months old (Fig 2), which does not mimic the phenotype of humans with the equivalent mutation in heterozygosis (Mencia et al. 2009; Modamio-Hoybjor et al. 2004).

**Figure 2.**
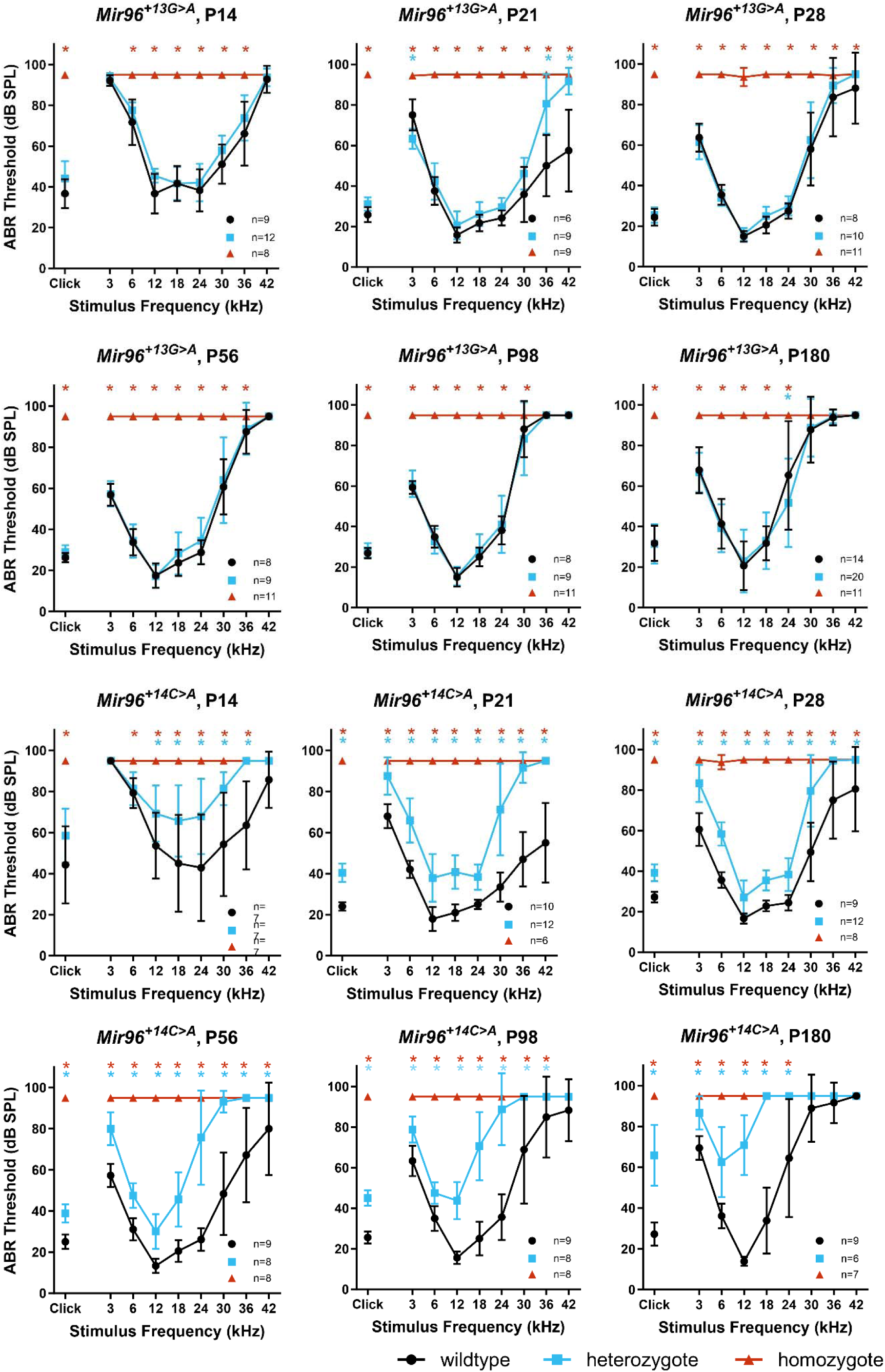
ABR thresholds of *Mir96^+13G>A^*and *Mir96^+14C>A^* mice at postnatal day (P)14, P21, P28, P56, P98 and P180. ABR measurements in response to click stimuli and tone pips ranging from 3 to 42 kHz were recorded from anaesthetised mice at different ages from P14 to P180. Homozygous mice for either mutation are shown as red triangles, heterozygotes as blue squares, and wildtypes as black circles. Each point in the plot shows the mean of the lowest stimulus level (threshold) at which a response is observed, +/-SD. Points at 95dB sound pressure level (SPL) indicate that there is no response up to the maximum sound level used. Both males and females were tested and are plotted together; n numbers are shown on each plot. Asterisks indicate significant differences (Bonferroni-corrected p < 0.05, mixed linear model pairwise comparison) between heterozygous mice and wildtypes (blue) or homozygous mice and wildtypes (red).

The mice were generated and maintained on the C57BL/6N genetic background. C57BL/6N mice are known to have age-related hearing loss, which is caused in part by the *Cdh23^ahl^* allele (Noben-Trauth, Zheng, and Johnson 2003). At 4 weeks of age, the high frequencies are affected, whereas the lower frequencies remain unaffected up to 6 months of age (Li and Borg 1991). We observed a similar pattern in wildtype mice from the *Mir96^+13G>A^* and *Mir96^+14C>A^* lines, which showed mild hearing loss at 24-42 kHz from 8 weeks old, but retained good low-frequency hearing sensitivity up to 6 months old (Fig 2).

### *Mir96^+13G>A^* heterozygotes have normal waveforms and retain normal hearing on a different genetic background and when subjected to noise

In order to further investigate the hearing of *Mir96^+13G>A^*heterozygotes, we first plotted their waveforms at 6 months old, and found no obvious difference when compared to the wildtype waveforms (Fig 3A). Mice were aged to 1 year old, but there was still no difference in the thresholds (Fig 3B).

**Figure 3.**
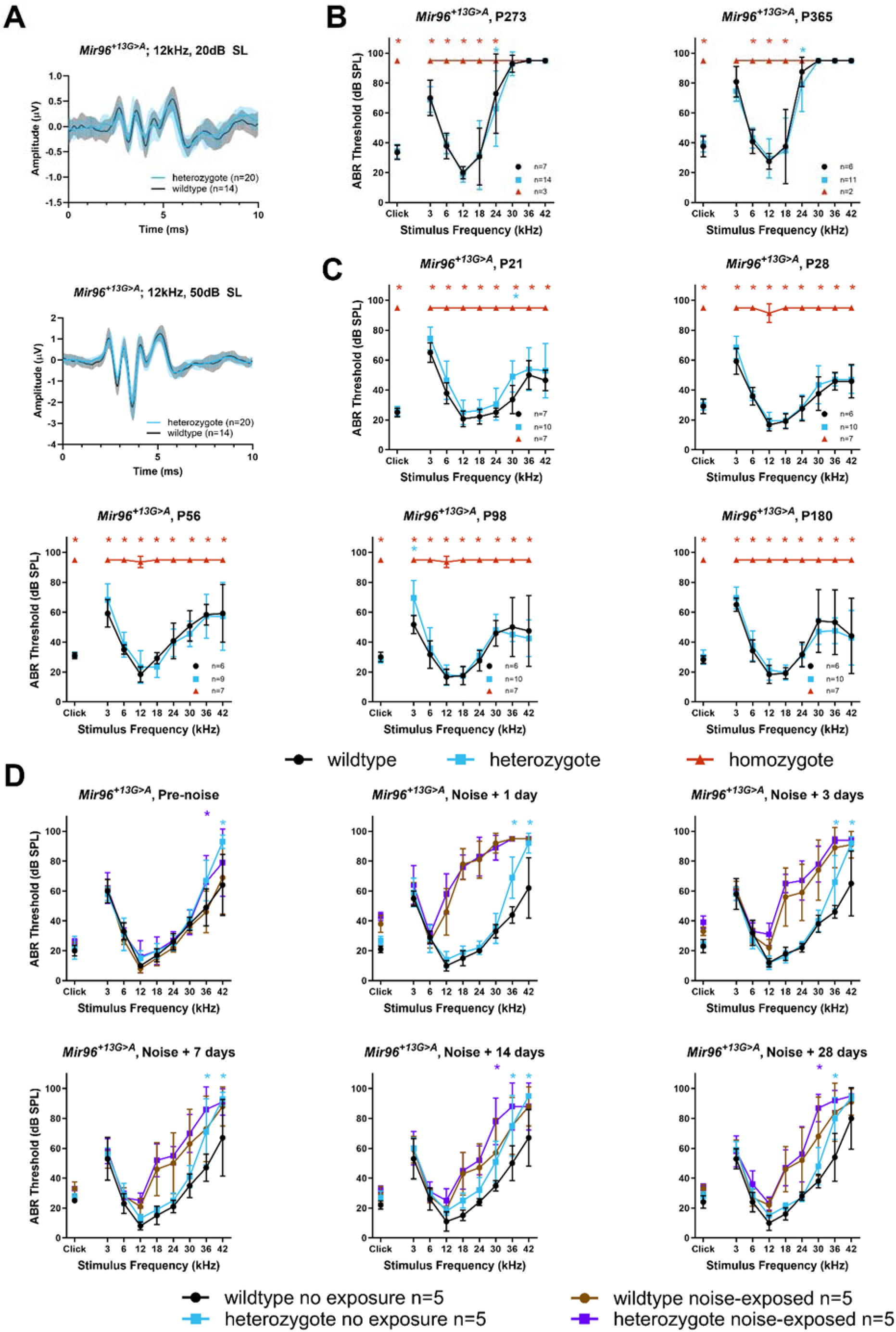
Further electrophysiological investigation of the *Mir96^+13G>A^* heterozygous mice. A) Mean ABR waveforms at 12 kHz, shown at 20 dB (top) and 50 dB (bottom) above threshold (sensation level, SL) ± standard deviation, at six months old. There is no obvious difference in waveform between *Mir96^+13G>A^* heterozygous (blue, n=20) and wildtype mice (black, n=14). B) Mean ABR thresholds from *Mir96^+13G>A^*homozygous (red triangles), heterozygous (blue squares) and wildtype (black circles) mice at 9 months and 1 year old. Males and females are plotted together; n numbers are shown on each plot. C) Mean ABR thresholds from *Mir96^+13G>A^*homozygous (red triangles), heterozygous (blue squares) and wildtype (black circles) mice on a mixed C3HeB/FeJ, C57BL/6N background at ages from 3 weeks to 6 months old. Males and females are plotted together; n numbers are shown on each plot. For B and C, asterisks indicate significant differences (Bonferroni-corrected p < 0.05, mixed linear model pairwise comparison) between heterozygous mice and wildtypes (blue) or homozygous mice and wildtypes (red). D) Mean ABR thresholds from *Mir96^+13G>A^*heterozygous and wildtype mice before and at multiple time points after noise exposure, showing similar recovery of thresholds in wildtype (brown circles, n=5) and heterozygous (purple squares, n=5) mice. Control wildtype (black circles, n=5) and heterozygous (blue squares, n=5) littermates went through the same set of ABR measurements and spent the same time in the noise exposure chamber, but without the noise. Males and females are plotted together. Asterisks indicate significant differences (Bonferroni-corrected p < 0.05, mixed linear model pairwise comparison) between unexposed wildtype and unexposed heterozygous mice (blue) or noise-exposed wildtype and noise-exposed heterozygous mice (purple). Error bars in all panels are standard deviation.

Next the hearing of wildtype, heterozygous and homozygous *Mir96^+13G>A^*mice on a mixed C57BL/6N, C3HeB/FeJ background was tested. The C3HeB/FeJ strain was selected for this outcross because it is known to show good hearing into old age and was the genetic background of the *Mir96^Dmdo^* mutant line previously studied (Lewis 2009). However, on the mixed background no difference was found between wildtypes and heterozygotes up to 6 months old, while homozygotes were still profoundly deaf at all ages tested (Fig 3C).

Finally, *Mir96^+13G>A^* heterozygote and wildtype mice were subjected to 100dB SPL, 8-16kHz noise for one hour to ask if the heterozygotes were more sensitive to noise damage than wildtypes. The noise-exposed mice exhibited a threshold shift which gradually recovered over the subsequent four weeks, but there was no obvious difference in thresholds or in threshold recovery rate between the wildtype and heterozygous mice (Fig 3D). The difference in the mean thresholds of noise-exposed heterozygotes and wildtypes at 30kHz 14 and 28 days after noise is statistically significant, but there is a lot of variability between individual mice, so we would not conclude that this is a biologically relevant difference.

While the *Mir96^+13G>A^* heterozygous non-exposed mice do appear to have worse thresholds than the non-exposed wildtypes, this is because these five wildtype mice have better high-frequency thresholds than the mice which went through the initial ABR tests (Fig 2).

### *Mir96^+13G>A^* and *Mir96^+14C>A^* homozygous mice have severely affected stereocilia bundles

Scanning electron microscopy (SEM) showed that homozygous mice of either mutation have severely affected stereocilia bundles at postnatal day 28 (P28). Stereocilia defects were seen in both OHCs and IHCs and were more severe in *Mir96^+14C>A^* than in the *Mir96^+13G>A^* mutants. We observe disorganised stereocilia that had lost their normal staircase arrangement, a fusion of stereocilia, and giant stereocilia, particularly in the IHCs (Fig 4), as well as missing stereocilia bundles. That phenotype becomes more striking at high frequency regions.

**Figure 4.**
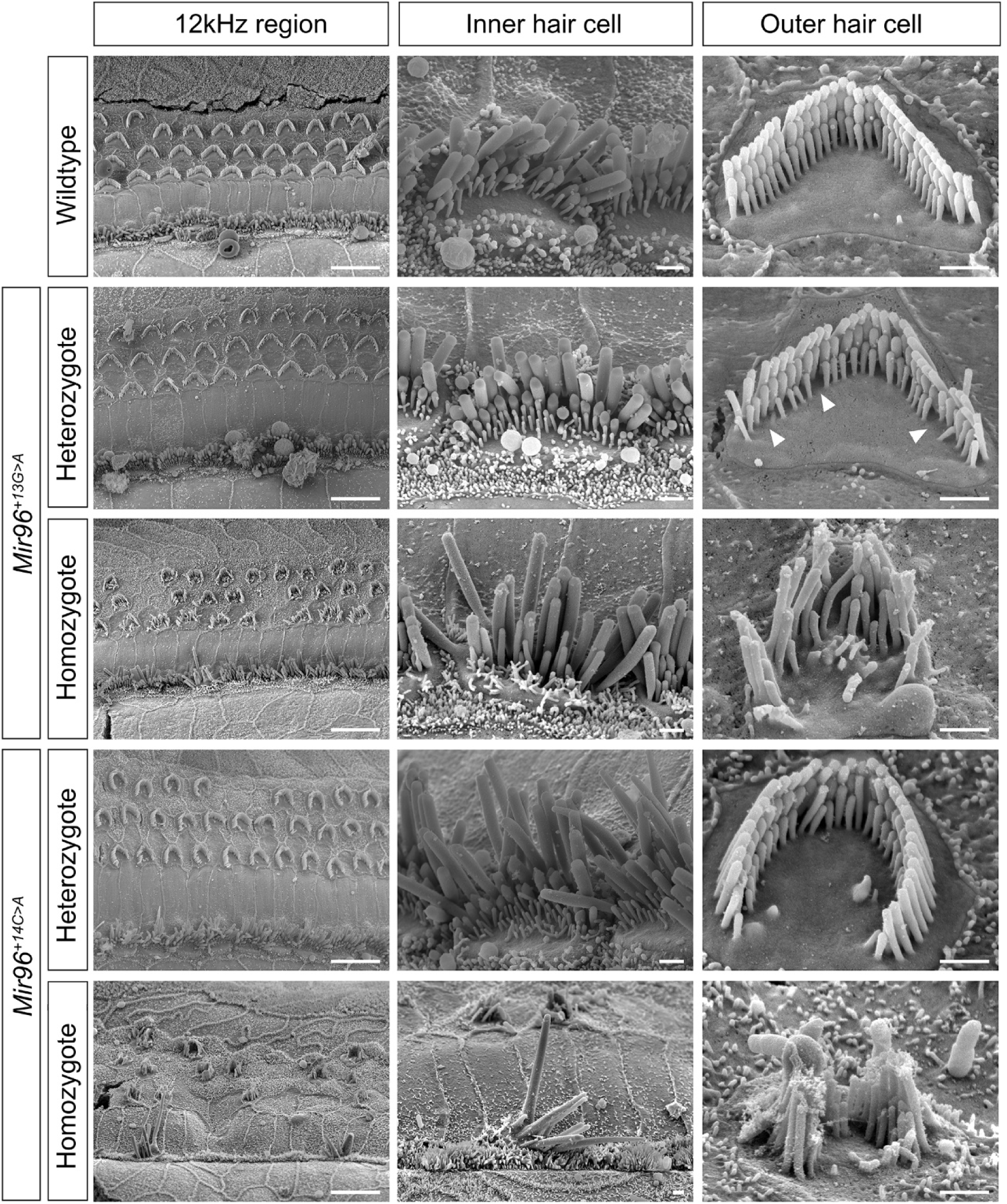
Scanning electron micrographs of *Mir96^+13G>A^* and *Mir96^+14C>A^*mice at 4 weeks old. Representative examples of wildtype, heterozygous and homozygous mice are shown. The images correspond to the 12 kHz best frequency region. For each panel, the left column shows a zoomed-out image with inner and outer hair cell rows. The middle column shows an inner hair cell close up, and the right column shows an outer hair cell close-up. *Mir96^+13G>A^* mice: wildtype (n=4), heterozygote (n=3), homozygote (n=3). *Mir96^+14C>A^* mice: wildtype (n=2), heterozygote (n=6), homozygote (n=3). Arrowheads point to the loss of stereocilia in the shortest row of the OHC bundles in *Mir96^+13G>A^* heterozygotes. Scale bar on left hand panels = 10 µm; scale bar for single hair cells = 1 µm.

In the heterozygotes, stereocilia damage is less severe than in the homozygotes. *Mir96^+13G>A^* heterozygous mice, despite having normal ABR thresholds, show loss of some stereocilia in the shortest row of the OHC bundles, which worsens at higher frequency regions. In *Mir96^+14C>A^* heterozygotes, which have mild hearing loss by P28, some OHC stereocilia bundles have a U-shape, instead of the typical V-shape observed in wildtypes; this rounding is more often seen towards the apical turn of the cochlea (Fig 4). In inner hair cells of *Mir96^+14C>A^* heterozygotes, some of the stereocilia appeared to taper towards their tips (Fig 4).

### *Mir96^+13G>A^* homozygous mice show a reduction in the number of IHC synapses

Confocal microscopy was used following immunolabelling of the pre- and post-synaptic markers CtBP2 and GluR2, respectively, to study the synapses at four weeks old (Fig 5A). *Mir96^+13G>A^* homozygotes were found to have a significant reduction in the number of ribbon synapses per inner hair cell (defined as colocalised pre- and postsynaptic labelling) compared to wildtypes at four weeks old (Fig 5B). There were no differences in heterozygotes compared to wildtypes. Furthermore, no significant differences were found in the number of synapses in *Mir96^+14C>A^* wildtypes, heterozygotes, and homozygotes (fig 5C).

**Figure 5.**
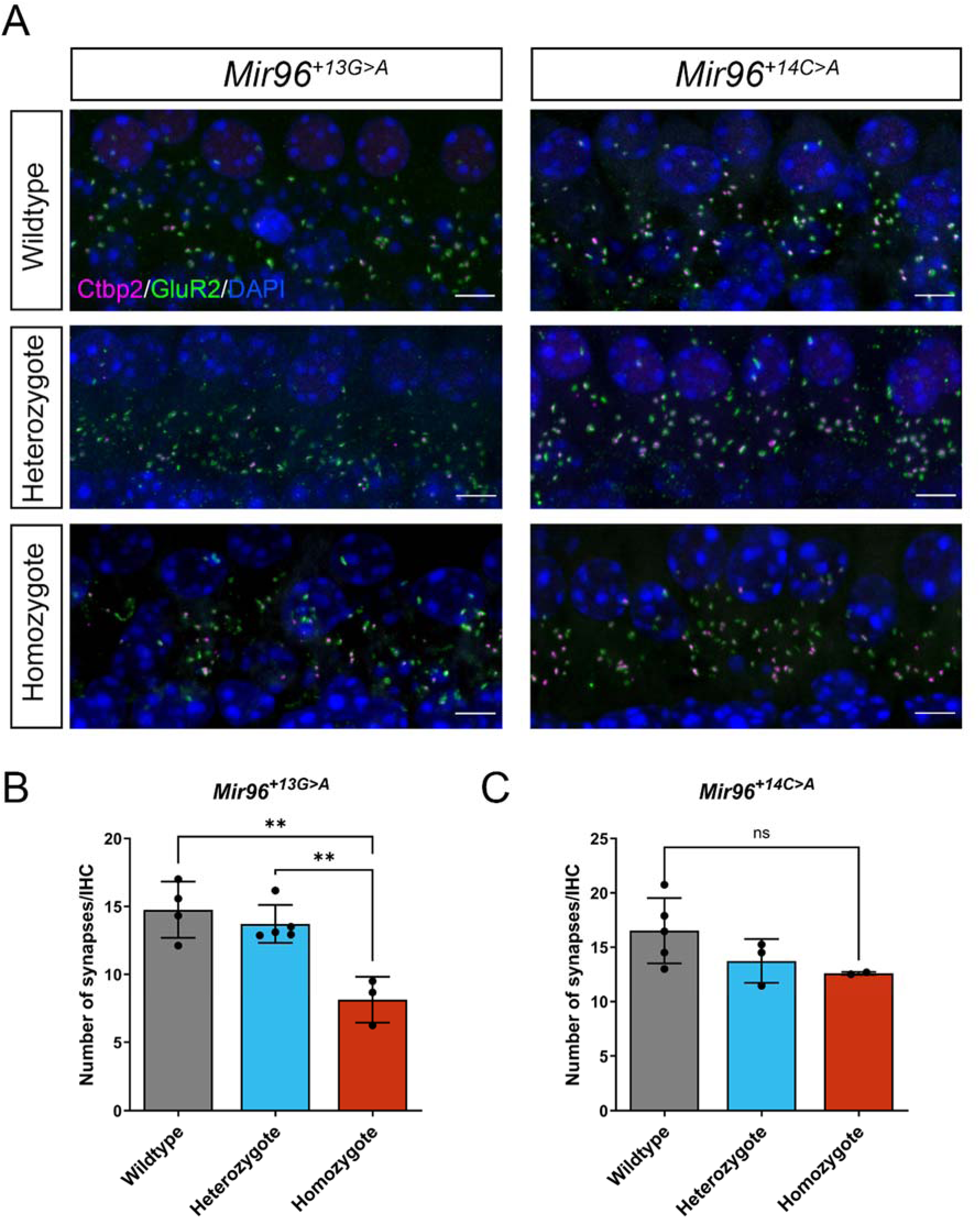
Analysis of synapses in *Mir96^+13G>A^* and *Mir96^+14C>A^* mutant mice at 4 weeks old. A) Confocal images of the whole-mount organ of Corti. Synapses were examined using an anti-CtBP2 antibody to mark pre-synaptic ribbons (pink) and an anti-GluR2 antibody to mark postsynaptic densities (green). Nuclei are shown in blue (DAPI). The images correspond to the cochlear region of 12 kHz best frequency. Scale bar=5µm. B, C) Quantification of ribbon synapses per IHC in *Mir96^+13G>A^* (B) and *Mir96^+14C>A^* (C). Confocal z-stacks were obtained with a z-step size of 0.25 µm and maximum intensity projection images were used for synapse counting. Colocalised pre and postsynaptic components were defined as a synapse. Synapses were counted and divided by the number of IHCs, determined by Myo7a staining (not shown in the images, only used for quantification purposes). All data are shown as mean ± SD and statistically analysed by one-way ANOVA with Tukey’s multiple comparisons test (** = P < 0.01). *Mir96^+13G>A^* mice: wildtype (n=4), heterozygote (n=5), homozygote (n=3); p=0.0016 (wildtype vs heterozygote adj. p=0.65; heterozygote vs homozygote adj. p=0.0041; wildtype vs homozygote adj. p=0.0018). *Mir96^+14C>A^* mice: wildtype (n= 5), heterozygote (n=3), homozygote (n=2); p=0.18.

### *Ocm* and *Slc26a5* are downregulated in *Mir96^+13G>A^* and *Mir96^+14C>A^* homozygotes

*Ocm* and *Slc26a5* are two genes that are strongly expressed in normal outer hair cells and both were strongly downregulated in *Mir96^Dmdo^* mice (Lewis et al. 2009) and in *Mir183/96^dko^* mice (Lewis et al. 2020). Therefore we measured expression levels of these two genes in the two new mutants studied here using qPCR on RNA from the organ of Corti at P4, and both genes were found to be significantly downregulated in *Mir96^+13G>A^* and *Mir96^+14C>A^*homozygotes (Fig 6).

**Figure 6.**
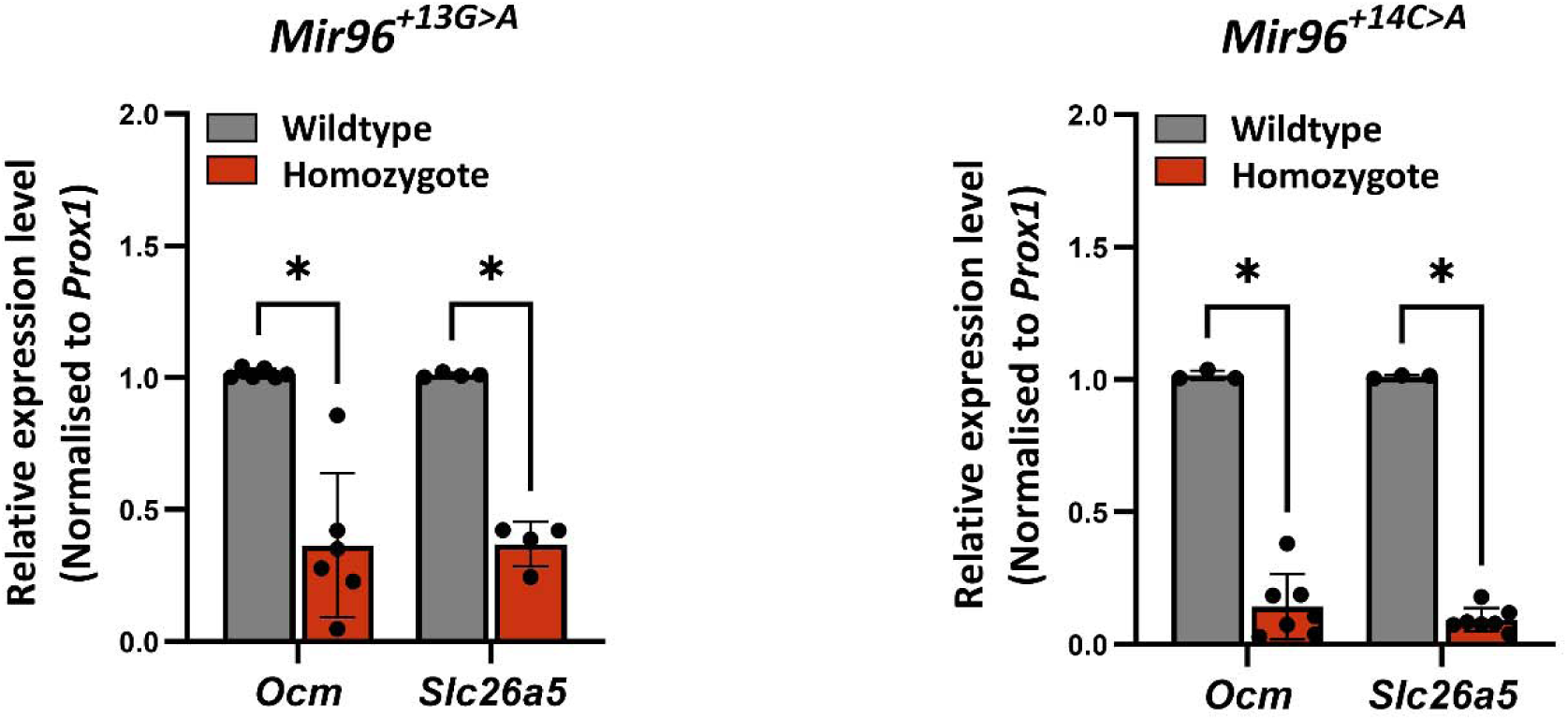
*Ocm* and *Slc26a5* are downregulated in *Mir96^+13G>A^* and *Mir96^+14C>A^*mutants. RT-qPCR was carried out on cDNA from the organ of Corti of P4 wildtype (grey) and homozygous (red) littermates to test gene expression changes. Mean expression levels were calculated from individual expression levels from each mouse and normalised to expression in a wildtype littermate. Relative expression levels were determined using the 2^-ΔΔct^ equation (Livak and Schmittgen 2001), using *Prox1* as an internal control for the amount of organ of Corti tissue present because *Ocm* and *Slc26a5* are specifically expressed in hair cells whereas *Prox1* is expressed in supporting cells of the organ of Corti (Bermingham-McDonogh et al. 2006). Error bars represent the standard deviation (Wilcoxon test; * = P < 0.05). *Mir96^+13G>A^ Ocm* = 6 wildtypes, 6 homozygotes, p=0.0022; *Slc26a5* n = 4 wildtypes, 4 homozygotes, p=0.029. *Mir96^+14C>A^ Ocm* = 3 wildtypes, 7 homozygotes, p=0.017; *Slc26a5* n = 3 wildtypes, 7 homozygotes, p=0.017. At least three technical replicates were used for each experiment.

### *Mir96^+14C>A^* homozygous mice have a larger number of differentially expressed genes than *Mir96^+13G>A^* homozygotes

In order to further explore the transcriptome of these mice, we carried out RNA-seq analysis of the organ of Corti of homozygotes and wildtype sex-matched littermates at P4. This age was chosen because earlier analyses had shown that all hair cells were still present at this early age, and we wanted to detect expression changes reflecting the different genotypes of the mice rather than due simply to reduced numbers of the cell type of interest (hair cells). Our results revealed that many genes are dysregulated in the organ of Corti of *Mir96^+13G>A^* and *Mir96^+14C>A^*homozygous mice, confirming that miR-96 controls a complex network of genes in the inner ear (Fig 7, Table S3). *Mir96^+13G>A^* homozygotes have 328 significantly differentially expressed genes (DEGs) (FDR < 0.05), 203 of which are upregulated and 125 downregulated. In *Mir96^+14C>A^* mutants, a total of 693 genes are significantly differentially expressed, with 369 upregulated and 324 downregulated. The much larger number of DEGs in *Mir96^+14C>A^* homozygous mice compared to *Mir96^+13G>A^* homozygotes correlates with the more severe structural phenotype observed in these mutants (Fig 4). The two mutants share only 124 DEGs misregulated in the same direction, indicating that transcriptomic changes due to each mutation are very different, even though both mutations are only one base apart in the *Mir96* sequence (Fig 7, Table S3).

**Figure 7.**
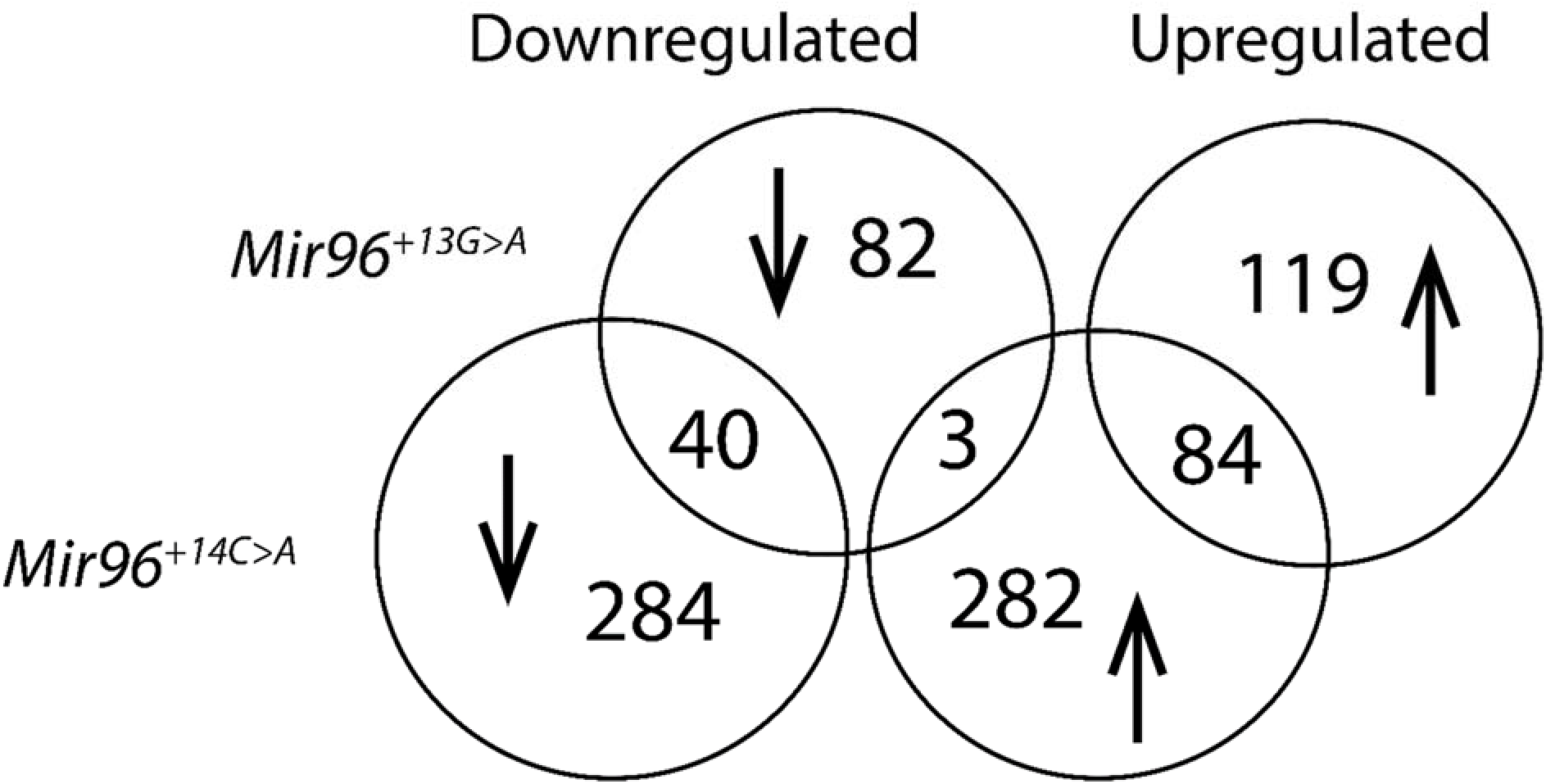
Comparison of up- and downregulated genes in *Mir96^+13G>A^* and *Mir96^+14C>A^*homozygotes. The majority of differentially expressed genes (DEGs) are not shared. Three genes are upregulated in *Mir96^+14C>A^* and downregulated in *Mir96^+13G>A^* homozygotes.

### Sylamer analysis identifies both loss of wildtype targets and gain of novel targets in each mutant

We used Sylamer (van Dongen, Abreu-Goodger, and Enright 2008) to assess the impact of the two mutations on the mRNA profile of both mutants. Analysis of all heptamers shows that the complementary heptamer to the miR-96 seed region (GTGCCAA, red line) is greatly enriched in the 3’UTRs of hundreds of genes upregulated in *Mir96^+13G>A^* and *Mir96^+14C>A^*homozygotes (Fig 8). This indicates that miR-96 normally represses a wide range of target genes, and when it is mutated, it is not able to repress its targets, which then become upregulated.

**Figure 8.**
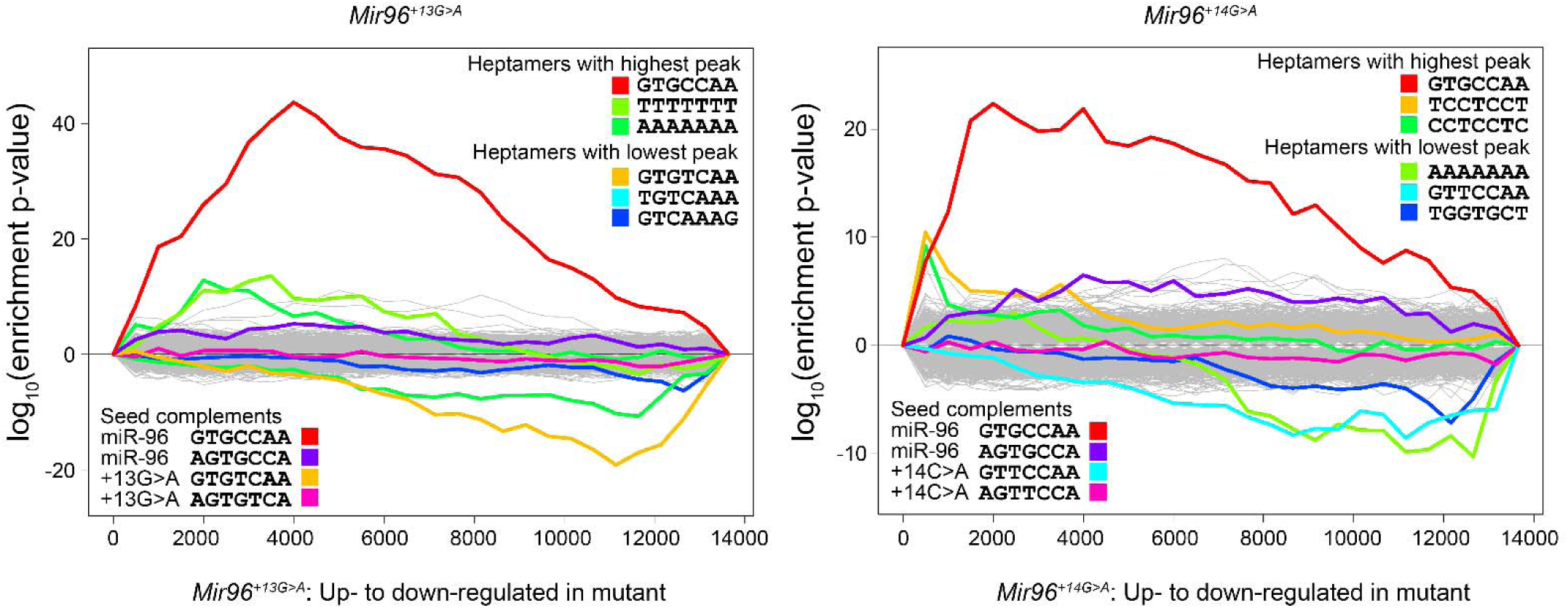
Sylamer enrichment landscape plots for *Mir96^+13G>A^* and *Mir96^+14C>A^* homozygous mice. The plots show enrichment and depletion of heptamers in the 3’UTRs of the differentially expressed genes (DEGs). The x-axis shows the list of DEGs for each mutant ranked from most upregulated (left) to most downregulated (right), based on the fold change (logFC). The Y axis shows the hypergeometric significance for the enrichment or depletion of heptamers in 3’UTRs in the leading parts of the gene list. Positive values indicate enrichment and negative values indicate depletion. The grey lines show the profiles of unenriched heptamers, while the coloured lines represent either heptamers that are complementary to the seed region of miR-96, or the lines with the highest and lowest peaks. Sylamer measures the enrichment of every possible heptamer in the 3’UTRs of the genes, in cumulative bins of 500 (x-axis), and generates a landscape plot. In this case, the main peaks in both mutants are from the wildtype seed (red) and show that it is enriched in the upregulated genes. The negative peaks on the right hand of each plot indicate that the downregulated genes are enriched in heptamers corresponding to the mutant miR-96. In the *Mir96^+13G>A^*graph, the yellow line corresponds to the mutant seed region, and in the *Mir96^+14C>A^* graph, the main peak of the mutant seed region is indicated in cyan. These negative peaks suggest that the mutant miR-96 is acquiring new targets.

We then asked whether mutations led to the silencing of potential acquired targets, that is, those genes containing in their 3’UTR binding sites complementary to the mutant seed region of miR-96. We observed that the heptamer complementary to the mutant miR-96 is enriched among the most downregulated genes, indicating that mutant miR-96 influences the expression of newly acquired target genes. The complementary heptamer to the *Mir96^+13G>A^* mutant seed region is GTG<u>T</u>CAA (Fig 8, yellow line), and the one complementary to the *Mir96^+14C>A^* mutant seed region is GT<u>T</u>CCAA (Fig 8, cyan line).

### The differentially expressed genes in *Mir96^+13G>A^* and *Mir96^+14C>A^* mutants are enriched in specific processes

We carried out Gene Set Enrichment Analysis (GSEA) (Mootha et al. 2003; Subramanian et al. 2005) on the transcriptomes from each mutant (Table S4), and visualised the results in Cytoscape. An enrichment map was generated for each mutant (Fig S1) (Reimand et al. 2019). From the *Mir96^+13G>A^*enrichment map (Fig S1A), we observed that a large number of DEGs were involved in synaptic activity, with terms like “presynaptic”, “dopamine transport” and “NMDA activation” being enriched, which correlates with the synaptic phenotype in this mutant (Fig 5). In the *Mir96^+14C>A^* network (Fig S1B), we observed gene sets involved in functions related to the cytoskeleton, extracellular matrix and adhesion molecules, which may be connected to the severe stereocilia degeneration observed in these mice (Fig 4).

We also analysed the significantly misregulated genes using Ingenuity Pathway Analysis (Qiagen, Germany), which identified canonical pathways with a significant overlap with the misregulated genes. There were 10 pathways with significant overlaps in the *Mir96^+13G>A^* data, including sensory processing of sound by outer and inner hair cells of the cochlea, and several pathways involved in Notch signalling (Table S5). From the *Mir96^+14C>A^*data, we obtained 45 significant pathways, which also included sensory processing of sound and Notch signalling, as well as degradation of the extracellular matrix, assembly of collagen fibrils, and integrin cell surface interactions (Table S5).

### Comparison of the misregulated genes with *Mir96^Dmdo^* and *Mir183/96^dko^* mutants

Our aim in investigating the transcriptome of these *Mir96* mutant mice was to determine candidate proteins with therapeutic potential. However, comparison of DEGs in *Mir96^+13G>A^*and *Mir96^+14C>A^* with previous data from *Mir96^Dmdo^*( (Lewis et al. 2016), Table S4) and *Mir183/96^dko^* (Lewis et al. 2020) (Fig 9) revealed only six genes shared between the four mutants. *Myo3a*, *Hspa2* and *St8sia3* are upregulated, while *Tmc1*, *Slc26a5* and *Ocm* are downregulated. Some other genes are differentially expressed in just two or three of the mutants, but most of them are differentially expressed in only a specific mutant, highlighting the transcriptomic differences obtained as a consequence of the different mutations and the different approaches. Furthermore, some genes are misregulated in two or more mutants but in different directions. For example, *Otof* is downregulated in *Mir96^+13G>A^*and *Mir96^+14C>A^* mutants, but it is upregulated in *Mir96^Dmdo^* mice.

**Figure 9.**
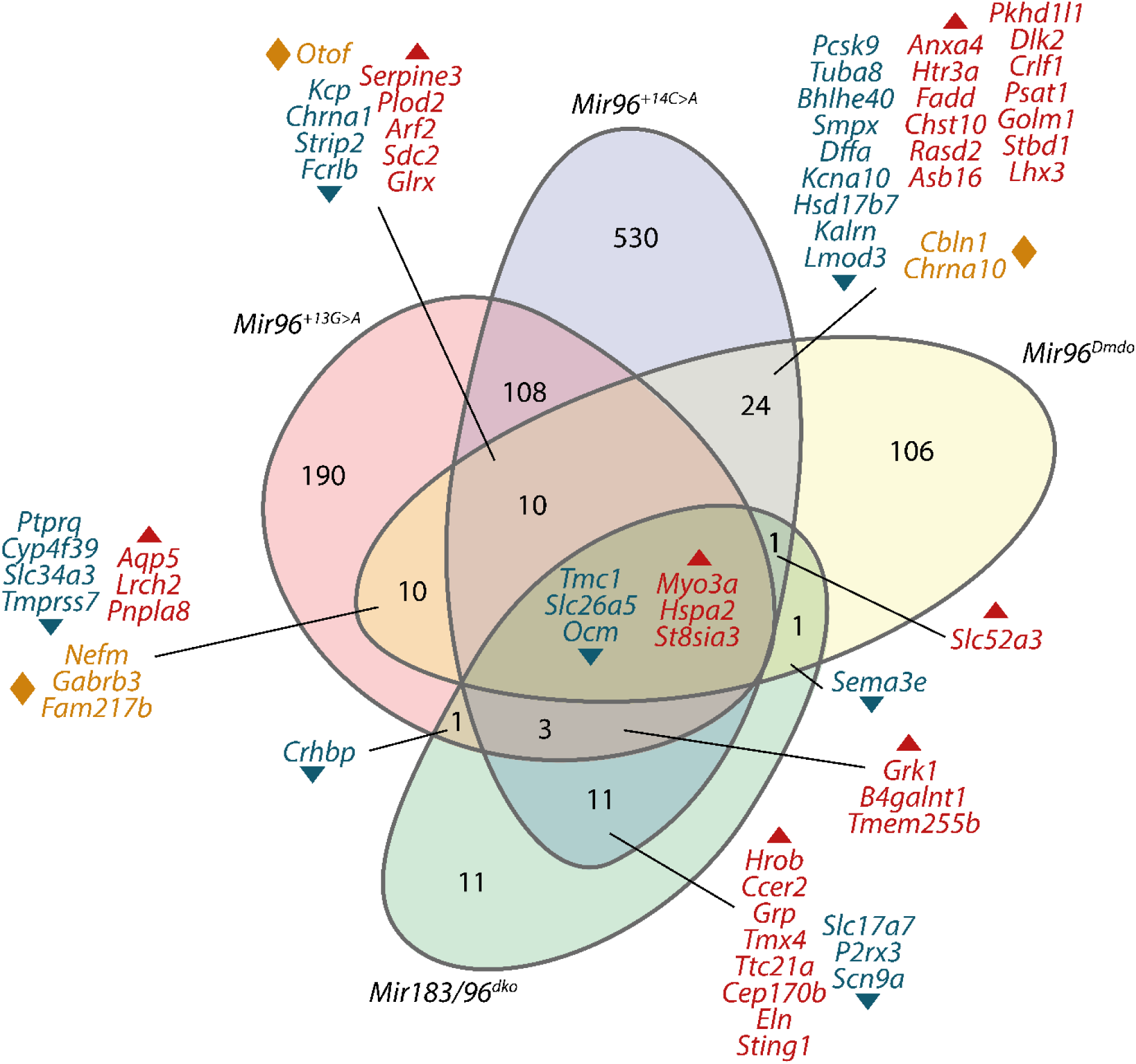
Comparison of the differentially expressed genes between four miR-96 mutants: *Mir96^+13G>A^*, *Mir96^+14C>A^*, *Mir96^Dmdo^* and *Mir183/96^dko^*. Downregulated genes are indicated in dark teal, upregulated genes in red, and genes that are upregulated in one mutant and downregulated in another are shown in orange.

### Identification of candidate miR-96 targets

Identifying targets of a master regulator is an important step in building a network which can then be assessed for therapeutic potential. We took three approaches to investigate potential direct targets of miR-96 in all four mutant mice.

First, we used Sylamer with different word lengths (6, 7 (as above) and 8) and identified the peak enrichment of the wildtype seed region closest to the start of the ranked genes (ranked from most upregulated to most downregulated). Genes ranking higher than this threshold (that is, to the left of this peak, shown by a vertical line in Fig S2) whose 3’UTR sequences contain matches to the miR-96 seed region are candidate targets (van Dongen, Abreu-Goodger, and Enright 2008). Using this approach, we identified peaks from all four mutant mice (Fig S2) and obtained 526 candidate genes for *Mir96^+13G>A^*, 355 for *Mir96^+14C>A^*, 386 for *Mir183/96^dko^*and 570 for *Mir96^Dmdo^*. There were 41 candidates shared between all four lists (Table S6), including one known deafness gene (*Eps8l2*, (Dahmani et al. 2015; Furness et al. 2013)).

Secondly, we used the gene set enrichment analysis to identify gene sets defined by a common regulator, usually identified by the presence of a binding motif close to their start sites. In the *Mir96^+13G>A^*analysis there were 15 known transcription factor gene sets in the upregulated genes, and none in the downregulated genes (Table S4). In the *Mir96^+14C>A^*analysis there were two transcription factor gene sets in the upregulated genes and 49 known transcription factor gene sets in the downregulated genes (Table S4). Of these transcription factors, Meis1, Mef2a and Jun (in the *Mir96^+13G>A^* gene sets) and Mafg, Mef2a, Foxf2, Gtf2a1, Nfkb1, Alx4, Mecom, Rreb1 and Mef2c (in the *Mir96^+14C>A^*gene sets) have wildtype miR-96 seed region matches in their 3’ UTRs and are thus potential direct targets (Table S6). We then performed the same GSEA analysis on the *Mir183/96^dko^* and *Mir96^Dmdo^* transcriptomes (Lewis et al. 2020; Lewis et al. 2009) (Table S7). There were only three known transcription factor gene sets identified from the *Mir183/96^dko^* transcriptome, and none of them had wildtype miR-96 seed region matches in their 3’ UTRs. In the *Mir96^Dmdo^* transcriptome, we found 40 known transcription factor gene sets, and 24 of the associated transcription factors had wildtype miR-96 seed region matches in their 3’UTRs (Mef2c, Meis1, Mafg, Sox5, Tbp, Creb1, Mtf1, Pbx1, Zeb1, Ar, Atf2, Atf3, Esr1, Esrra, Foxo4, Hjurp, Jun, Mecom, Mef2a, Pax7, Phf21a, Runx1, Sp1, Vsx1, Zbtb14, Table S6). The only candidate target transcription factor shared between three mutant transcriptomes is Mef2a, which has previously been found to be expressed in spiral ganglion neurons, inner hair cells and the supporting cells of the organ of Corti at P15 (Sanchez-Calderon et al. 2010).

Thirdly, we used Ingenuity Pathway Analysis to predict upstream regulators (Kramer et al. 2014) based on the significantly misregulated genes in each of the four mutant miR-96 transcriptomes. From the list of potential upstream regulators, we selected those genes which were predicted to have higher activity in homozygotes and had wildtype miR-96 seed region matches in their 3’ UTRs. Any genes matching this description which were known to be downregulated in the transcriptome data were excluded. We obtained 24 candidates for *Mir96^+13G>A^*, 27 for *Mir96^+14C>A^*, 14 for *Mir96^Dmdo^* and 7 for *Mir183/96^dko^*. Only three candidate targets were shared; *Neurod4*, *Lrig1* and *Cdh1* were candidate targets in both the *Mir96^+13G>A^* and the *Mir96^+14C>A^* transcriptomes (Table S6).

Only two proteins were identified as candidates by more than one approach; Pax7 and Atf3, found in both the GSEA and IPA analyses. There were no overlaps between either the GSEA or the IPA target lists and the Sylamer targets.

### Identifying candidate therapeutics from whole transcriptome data

Because we were unable to identify a shortlist of candidate target genes or proteins to assess for therapeutic potential, we chose instead to use the whole transcriptome, comparing it to known drug profiles from DrugMatrix, which contains the results from thousands of experiments treating rats or rat cell lines with drugs (https://ntp.niehs.nih.gov/data/drugmatrix). This was carried out using the SPIED platform (www.spied.org.uk (Williams 2012, 2013)). In order to focus on the most important differentially expressed genes and to exclude, as much as possible, genes misregulated because they are novel targets of a specific mutant miR-96 seed, we looked at genes with consistent differential expression between the *Mir96^+13G>A^*and the *Mir96^+14C>A^* transcriptomes (7944 genes in total, Table S8). We did not include the other two transcriptomes because they were carried out on different platforms and with different methods, which could result in exclusion of genes for reasons unrelated to their biological relevance (for example, *Ccer2*, which is downregulated in the *Mir183/96^dko^*and the *Mir96^+14C>A^* transcriptomes, was not present on the microarray used for the *Mir96^Dmdo^* study).

SPIED outputs 100 profiles by default, ranked from the most similar to the input profile to the most dissimilar, and the drugs with the most dissimilar profile are potential therapeutics because they lead to a complementary change in transcription patterns across the 7944 genes tested. Among the anti-correlated drug profiles compared with the profiles from *Mir96^+13G>A^* and *Mir96^+14C>A^* (Table S8) is amitriptyline, a tricyclic antidepressant which has previously been reported to improve recovery after noise-induced hearing loss in guinea-pigs (Shibata et al. 2007). Amitriptyline was also identified in the SPIED analysis of the *Mir183/96^dko^* transcriptome, which should not include novel targets since the mutant allele is a deletion of *Mir96* (Table S8). It is known to cross the blood-brain barrier (Coudore et al. 1994), which suggests it may also be able to cross the blood-labyrinth barrier. We therefore considered it a good candidate for testing in *Mir96* mutant mice.

### Amitriptyline delays hearing loss in *Mir96^+14C>A^* heterozygotes

We first tested amitriptyline in wildtype C57BL/6N mice to verify that it did not affect hearing, and found that at 4 weeks old there was no difference between wildtype mice drinking water with saccharin and wildtype mice drinking water with saccharin and amitriptyline (Fig S3). We then tested *Mir96^+14C>A^*heterozygotes and homozygotes, and wildtype littermates, given either saccharin alone, saccharin and 200µg/ml amitriptyline, or saccharin and 400µg/ml amitriptyline. We observed progressive hearing loss in the heterozygotes drinking amitriptyline, but it was significantly delayed at high frequencies between 24 and 36 kHz compared to the heterozygotes drinking only saccharin, most visibly at 30kHz at 4 weeks old (Fig 10). No improvement was seen in the homozygotes drinking amitriptyline, and increasing the dose of amitriptyline to 400µg/ml made no difference to the hearing impairment in either heterozygotes or homozygotes (Fig 10).

**Figure 10.**
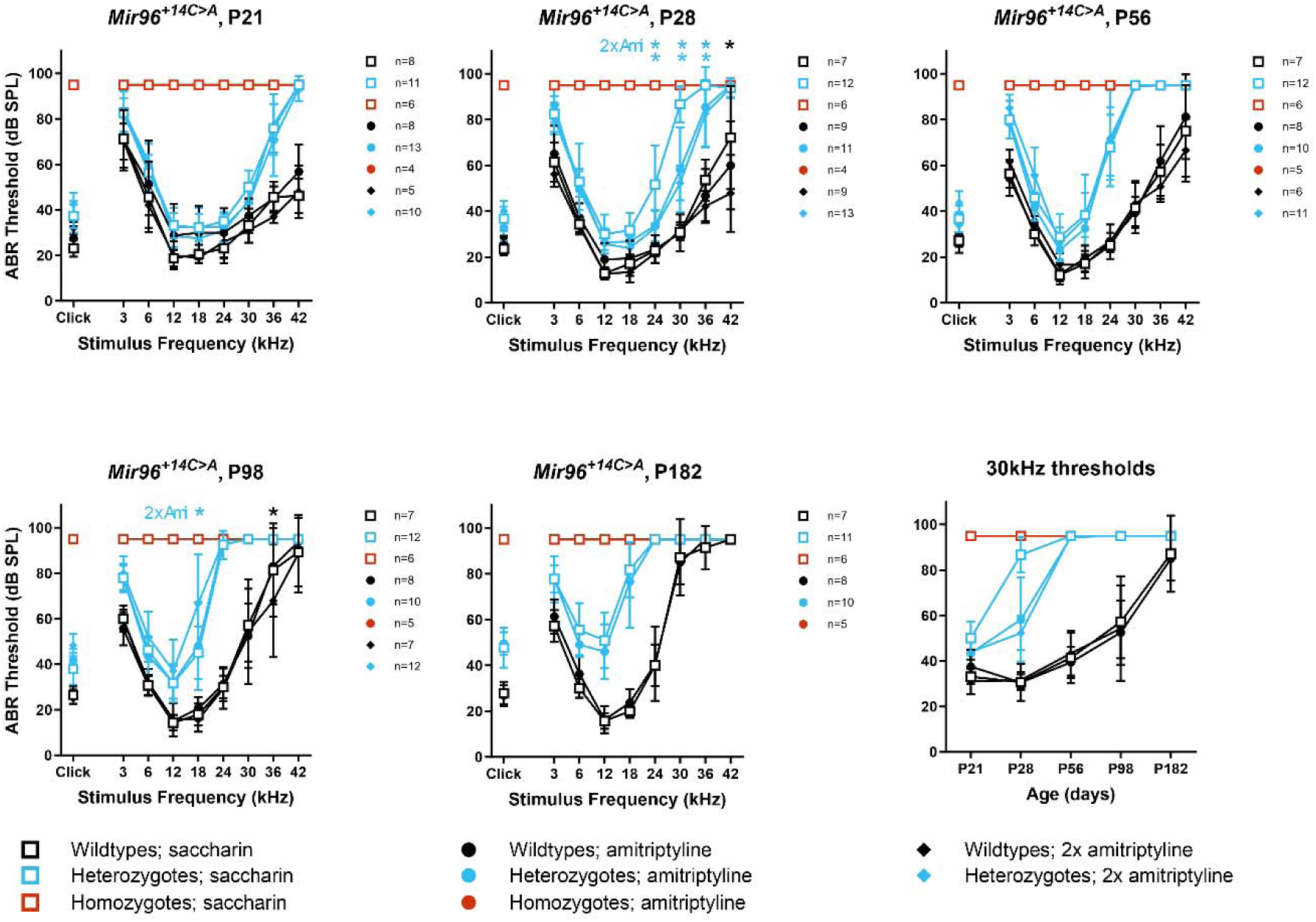
Amitriptyline delays the progression of hearing loss in *Mir96^+14C>A^* heterozygotes. Mean ABR thresholds from *Mir96^+14C>A^* heterozygous mice and wildtype littermates drinking either saccharin (open squares), 200µg/ml amitriptyline plus saccharin (circles), or 400µg/ml amitriptyline plus saccharin (diamonds) at ages from 21 days to six months. Asterisks indicate significant differences (Bonferroni-corrected p < 0.05, mixed linear model pairwise comparison) between mice drinking either amitriptyline or double-dose amitriptyline (2x Ami) compared to mice of the same genotype drinking saccharin. Significant differences between heterozygotes on differing dosages are marked in blue, significant differences between wildtypes on different dosages are marked in black. The final plot shows mean thresholds at 30kHz plotted against time. Error bars are standard deviation.

## Discussion

### Comparison with human phenotype and relevance to human *MIR96* mutations

The more severe phenotype observed in mutant mice with a point mutation in the seed region of miR-96 (*Mir96^Dmdo^*, *Mir96^+13G>A^*and *Mir96^+14C>A^*) compared to that of mice lacking miR-96 (*Mir183/Mir96^dko^*) suggests that the gain of novel targets plays an important role in the phenotype caused by miR-96 mutations. This has significant implications for the development of therapies. Since we know that the *Mir183/96^dko^* heterozygotes have normal hearing, one copy of wildtype miR-96 is enough for normal hearing function. Therefore, the use of a silencing RNA to target only the mutant copy of miR-96 may be enough to allow normal hearing.

While the *Mir96^+14C>A^* heterozygotes mimic the phenotype observed in the family with the equivalent mutation, the *Mir96^+13G>A^* heterozygotes escape deafness while the same mutation causes progressive hearing loss in humans (Mencia et al. 2009; Modamio-Hoybjor et al. 2004). *Mir96^+13G>A^* heterozygotes do not show raised thresholds up to 1 year old (Fig 2), in contrast to the progressive hearing loss observed in adulthood in the human counterparts. We tested these mice on a different genetic background and exposed them to noise, but found no differences in heterozygotes compared with wildtypes (Fig 3).

Therefore, we suggest that the normal hearing observed in *Mir96^+13G>A^* heterozygous mice compared to the progressive hearing loss observed in humans carrying the same mutation may be due to the repression of different novel target genes in humans and mice. One potential candidate is *RAB11A*, a gene which in humans has three matches to the *Mir96^+13G>A^* mutant seed region in its 3’UTR. There are no matches to the wildtype seed region, and no matches in the mouse *Rab11a* 3’UTR to either mutant or wildtype seed regions. This gene has recently been reported to be required for correct development of the stereocilia bundle in mice (Knapp et al. 2023). Identification of this and other candidates could be tested in future studies in cultured cells.

### The auditory phenotype of *Mir96^+14C>A^* mice is more severe than that of *Mir96^+13G>A^* mice but less severe than that of *Mir96^Dmdo^*mutants

Mice heterozygous for the *Mir96^+14C>A^* point mutation exhibit progressive hearing loss from four weeks old, while *Mir96^+13G>A^*heterozygotes have normal ABR thresholds (Fig 2).

*Mir96^Dmdo^* heterozygous mice (which carry the +15A>U point mutation) exhibit early-onset rapidly progressive hearing loss, even at two weeks old (Kuhn et al. 2011). Initially, it was thought that the *Mir96^Dmdo^*phenotype was due to haploinsufficiency rather than the acquisition of novel targets because humans heterozygous for different point mutations also have progressive hearing loss (Lewis et al. 2009; Mencia et al. 2009). However, deletion of miR-183 and miR-96 (*Mir183/96^dko^*) (Lewis et al. 2020) or the whole miR-186/96/182 cluster (Fan et al. 2017; Geng et al. 2018) does not lead to auditory dysfunction in heterozygosis.

Therefore, the phenotype observed in the *Mir96^Dmdo^* and *Mir96^+14C>A^* heterozygotes is likely to be partly due to the acquisition of new targets by the mutant miRNA.

Mice homozygous for all the *Mir96* alleles reported so far show no ABR responses at all ages tested (Table 1). This indicates that the abnormal targeting pattern of miR-96 in mutants affects the development of the hair cells, as previously described (Kuhn et al. 2011).

**Table 1.** Comparison of the phenotype in the three *Mir96* mutant mice carrying point

|  | Genotype | *Mir96*<sup>Dmdo</sup> | *Mir96*<sup>+14C>A</sup> | *Mir96*<sup>+13G>A</sup> | *Mir183/96*<sup>dko</sup> |
| --- | --- | --- | --- | --- | --- |
| **Genetic background** | **All the genotypes** | C3HeB/FeJ | C57BL/6N | C57BL/6N | C57BL/6N |
| **Auditory brainstem responses** | **Heterozygotes** | Early onset, rapidly progressive hearing loss | Progressive hearing loss | Normal hearing | Normal hearing |
|  | **Homozygotes** | Profound deafness | Profound deafness | Profound deafness | Profound deafness |
| **Surface of the organ of Corti** | **Heterozygotes** | Rounded bundles and disorganised stereocilia | Rounded OHC stereocilia bundles | Loss of stereocilia in the innermost row of the OHCs | OHCs have rounded stereocilia bundles. IHCs appear normal. |
|  | **Homozygotes** | Most hair cells have degenerated by 4 weeks old | Most bundles are missing | Severely affected stereocilia bundles | Severely affected IHCs and OHCs stereocilia bundles |
| **Synapses** | **Heterozygotes** | NA | Normal | Normal | Normal |
|  | **Homozygotes** | Immature IHC ribbon synapses | Normal | Significantly reduced at 4 weeks old | Significantly reduced at 4 weeks old |
| **Transcriptomics, number of DEGs (method)** | **Heterozygotes** | NA | NA | NA | NA |
|  | **Homozygotes** | 86 (Microarray) | 693 (RNA-seq) | 328 (RNA-seq) | 34 (RNA-seq) |

### Stereocilia bundles and ribbon synapses in *Mir96^+13G>A^* and *Mir96^+14C>A^* mice

Similar to the physiological phenotype, the structural phenotype of *Mir96^+13G>A^* is much less severe than that of *Mir96^+14C>A^* and *Mir96^Dmdo^*. Both inner and outer hair cells of *Mir96^+13G>A^* homozygotes show degenerative changes at four weeks old, but this phenotype is more severe in *Mir96^+14C>A^* homozygotes, where the stereocilia bundles are more severely affected by four weeks old (Fig 4), and even more severe in *Mir96^Dmdo^* homozygous mice, with very few stereocilia bundles visible in the organ of Corti at 4 weeks postnatal (Lewis et al. 2009). In *Mir183/96^dko^* homozygous mice, hair cells are also severely affected at 4 weeks of age, with many hair bundles missing entirely. Where present, the stereocilia bundles of both OHCs and IHCs show splaying and fusion (Lewis et al. 2020) (Table 1).

In *Mir96^+13G>A^* heterozygotes, a loss of some stereocilia in the shortest row of the OHC bundles was observed at 4 weeks old, when ABR thresholds were normal (Fig 4). Several cases of cochlear defects associated with normal ABR thresholds have been reported. For example, synaptic damage in the inner ear, also referred to as cochlear synaptopathy, can produce difficulties in understanding hearing speech in noisy environments without an increase in ABR thresholds (also known as hidden hearing loss) both after noise exposure (Kujawa and Liberman 2009) and ageing (Sergeyenko et al. 2013). Hidden hearing loss has also been linked to auditory nerve myelination defects (Wan and Corfas 2017) and auditory nerve dysfunction (Reijntjes et al. 2019). Furthermore, it has been reported that stereocilia tip tapering can also be associated with normal ABR thresholds (Ingham et al. 2021). Our observations suggest that the loss of stereocilia in the shortest row of the OHC bundles is an additional cochlear defect that can be hidden behind normal ABR thresholds.

Similarly to *Mir96^+14C>A^* heterozygous mice, *Mir96^Dmdo^* and *Mir183/96^dko^* heterozygotes have rounded OHC stereocilia bundles (Fig 4) (Lewis et al. 2020; Lewis et al. 2009). In *Mir96^+14C>A^* and *Mir96^Dmdo^* heterozygotes, the structural phenotype correlates with the progressive hearing loss observed in these mutants. However, *Mir183/96^dko^* heterozygotes have normal hearing.

In *Mir96^+13G>A^* homozygotes we found significantly fewer colocalised pre- and postsynaptic densities per IHC, indicating synaptic defects (Fig 5, Table 1). This was not observed in *Mir96^+14C>A^* homozygotes, suggesting that the mechanism of hearing loss caused by the two different mutations in *Mir96* is different. A reduced number of ribbon synapses per IHC was also observed in *Mir183/96^dko^* homozygotes (Lewis et al. 2020), and *Mir96^Dmdo^* homozygotes exhibit immature IHC ribbon shapes and disorganised innervation (Kuhn et al. 2011), although in this last mutant the synapses were not quantified in the same way so cannot be directly compared.

### Transcriptomic changes in *Mir96^+13G>A^* and *Mir96^+14C>A^* mutants

We have previously shown that many genes are dysregulated in *Mir96^Dmdo^* mice as a result of the point mutation affecting the seed region, indicating that miR-96 controls a complex network of genes in the inner ear (Lewis et al. 2016; Lewis et al. 2009). In the present study, we found 328 differentially expressed genes in *Mir96^+13G>A^* homozygotes and 693 in *Mir96^+14C>A^* homozygotes. RNA-seq of *Mir183/96^dko^* homozygotes revealed only 34 DEGs (Lewis et al. 2020) and microarray performed in *Mir96^Dmdo^* revealed 86 DEGs (Lewis et al. 2009). The milder effects observed in the knockout allele compared with point mutations in *Mir96* indicates that the gain of novel targets can play an important role in the phenotype caused by a microRNA mutation, as suggested by the Sylamer analysis (Fig 8). We conclude that the differentially expressed genes in *Mir96^+13G>A^*and *Mir96^+14C>A^* mutant mice are related to specific pathways and cellular processes that may be responsible for the phenotypes observed by SEM and confocal microscopy.

Many DEGs (124) were shared between *Mir96^+13G>A^* homozygotes and *Mir96^+14C>A^* homozygotes (Figure 7, Table S3), but only 6 were shared between all four *Mir96* mutants (Fig 9). This may in part be due to the different platforms used; the *Mir96^Dmdo^* data comes from a microarray and not RNA-Seq, and many genes were not represented on the microarray. It is also worth noting that *Mir96^Dmdo^* mice are on a C3HeB/FeJ genetic background, while *Mir96^+13G>A^*, *Mir96^+14C>A^*and *Mir183/96^dko^* mice are on a C57BL/6N background, which could explain some of the differences observed between the different mutants (Table 1). However, it is likely that much of the difference is due to the different mutations (three point mutations which affect targeting, and one knockout which includes the neighbouring *Mir183* gene).

### Identifying wildtype miR-96 targets

We made use of three different methods to identify potential targets of miR-96, but did not find any consistently identified candidates. It is possible that miR-96 operates mainly through mild downregulation of many direct targets, rather than strongly downregulating one or two, and thus it would be hard to identify individual target genes. One limitation of this study is that we carried out bulk RNA-seq of the whole organ of Corti, while miR-96 is only expressed in the hair cells. Therefore, if a target is highly expressed in the non-sensory epithelial cells, any difference in expression between *Mir96* wildtypes and homozygotes as a result of the repression in the hair cells might not be detectable. This might also explain why some genes that are known direct targets of miR-96, such as *Zeb1*, *Foxo1*, and *Nr3c1*, are not significantly misregulated in our transcriptomic data. Single-cell RNA sequencing (scRNA-seq) would be required to approach this problem.

It is also possible that miR-96 is regulating different genes in the inner and outer hair cells because the regulation is dependent on the mRNA molecules being expressed on a specific cell type. For instance, *Ocm* and *Slc26a5*, two of the most significantly downregulated genes, are predominantly expressed in outer hair cells. This would further complicate target identification.

### Gain of novel targets of mutant miR-96

The *Mir183/96* knockout mouse mutant (*Mir183/96^dko^*) provides a good comparison to study only the effects of the loss of normal miR-96 targets (since *Mir96* and *Mir183* are very closely linked, it was not possible to target either gene alone (Prosser et al. 2011)).

The lower number of DEGs obtained in the mice with the null allele (*Mir183/96^dko^*) compared to the mutants with a point mutation in the seed region of miR-96 suggests that the gain of novel target mRNAs is important for the phenotype of these mutants. Sylamer analyses (van Dongen, Abreu-Goodger, and Enright 2008) (Fig 8) supported that some of the wildtype miR-96 targets were upregulated in both mutants. However, there was also a marked negative peak for heptamers matching the mutant seed regions (Fig 8), meaning that some of the significantly downregulated genes bore matches to the mutant seed region in their 3’UTRs, and their downregulation is likely due to the acquisition of new targets by the mutant microRNA. The differences between the four *Mir96* mutants (Table 1) clearly demonstrates that the gain of novel targets plays an important role in the phenotype resulting from a point mutation in *Mir96*.

### Pharmacological interventions to maintain hearing: a proof of concept

Rather than focus on a single target, because our analyses suggest that miR-96 operates through a multitude of target genes, we used the whole transcriptome to identify drugs which have the opposite effect on gene expression. We chose amitriptyline to test, and found that it delays the progression of hearing impairment in *Mir96^+14C>A^* heterozygotes (Fig 10). This is a proof of concept rather than a suggestion that amitriptyline be used as a treatment for humans carrying the *MIR96^+14C>A^* mutation, for several reasons. First, the dosage in mice (by body weight) was far higher than the standard amitriptyline dose used in humans; second, amitriptyline can have multiple unpleasant side effects; and third, the delay in progression of hearing loss was only temporary. However, this shows that it is possible to intervene pharmaceutically to delay hearing loss caused by a genetic defect.

Currently there are no therapeutics for treating progressive hearing loss, so this is an important proof of principle. It is also worth noting that we were able to find a candidate therapeutic which had an effect even without a full understanding of the complex regulatory network controlled by miR-96, and this approach may generalise to other conditions and diseases caused by mutations in regulatory molecules.

## Acknowledgements

We thank Flavia Davidhi, Jack Blackburn, Susana Caetano for technical support, Neil Ingham for maintaining the auditory physiology facility, and the King’s College London Centre for Ultrastructural Imaging for maintaining the electron microscopy facility.

## Data availability

Transcriptome data from the two mouse mutants reported here are publicly available from ArrayExpress, accession number E-MTAB-13772 (https://www.ebi.ac.uk/biostudies/arrayexpress/studies/E-MTAB-13772).

