## Supplementary Figures 1-3 for "Pathological mechanisms and candidate therapeutic approaches in the hearing loss of mice carrying human *MIR96* mutations"

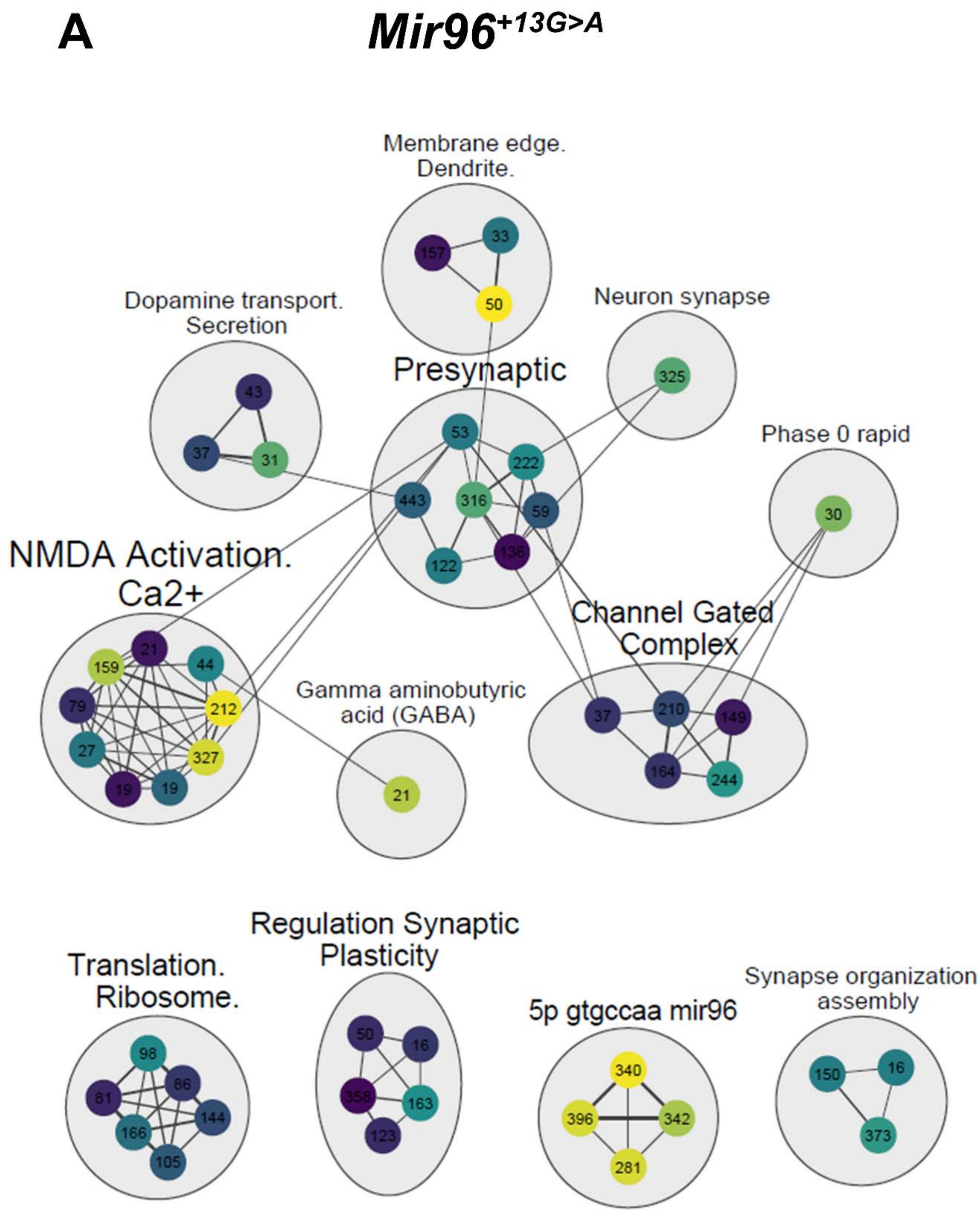

**B** ***Mir96***$^{+14C>A}$

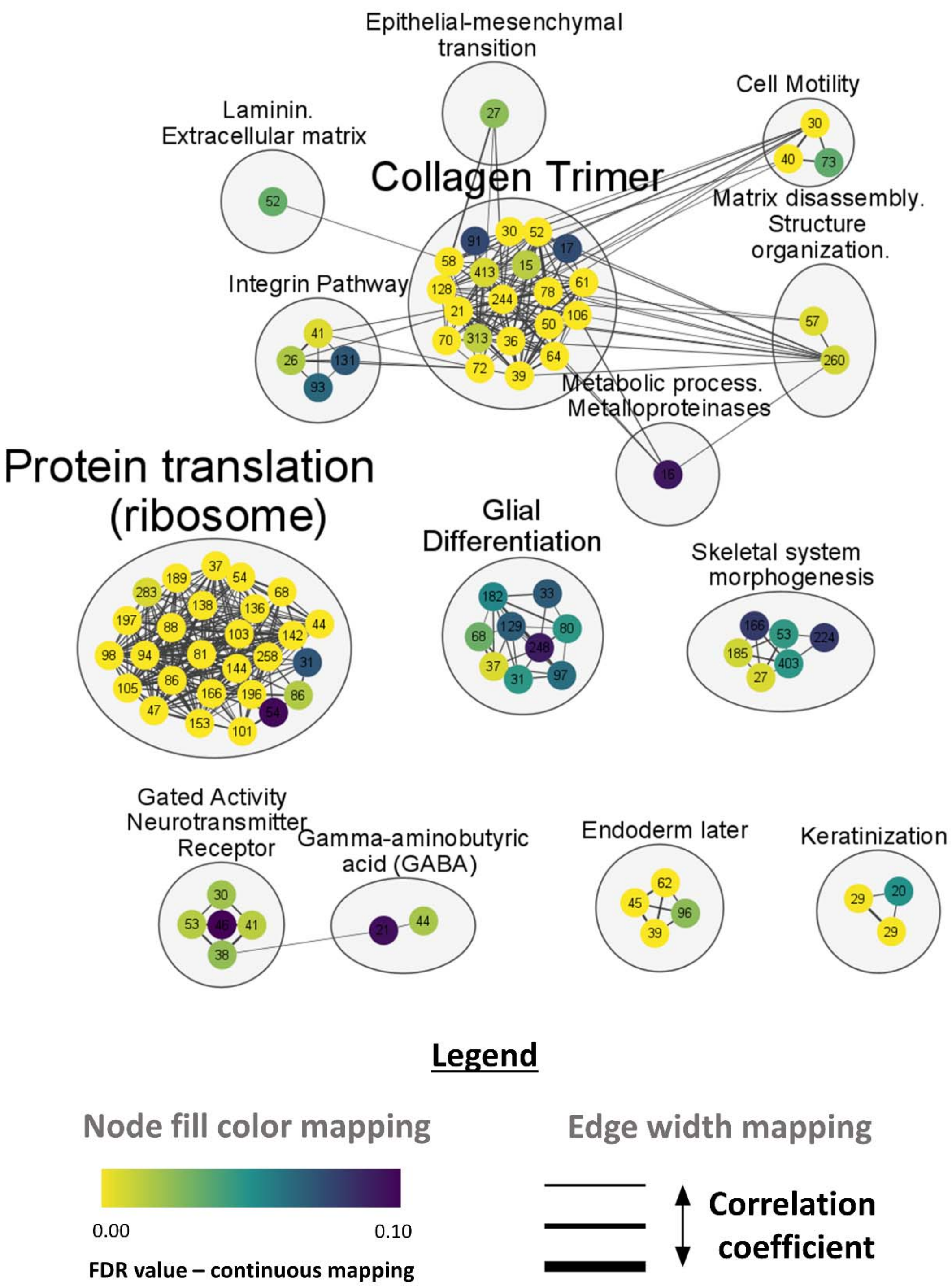

Epithelial-mesenchymal transition
Cell Motility
Laminin. Extracellular matrix
Collagen Trimer
Matrix disassembly. Structure organization.
Integrin Pathway
Metabolic process. Metalloproteinases
Protein translation (ribosome)
Glial Differentiation
Skeletal system morphogenesis
Gated Activity Neurotransmitter Receptor
Gamma-aminobutyric acid (GABA)
Endoderm later
Keratinization
Legend
Node fill color mapping
0.00
0.10
FDR value – continuous mapping
Edge width mapping
Correlation coefficient

**Figure S1. Enrichment map visualization in Cytoscape.** A) Enrichment map based on GSEA analysis of the *Mir96*$^{+13G>A}$ mutant transcriptome. B) Enrichment map based on GSEA analysis of the *Mir96*$^{+14C>A}$ mutant transcriptome. Each map contains gene sets (nodes) interconnected by the similarity between them (edges). The colour of the nodes is based on the FDR, with a continuous colour scheme from yellow (FDR=0) to purple (FDR=0.1). In other words, the nodes in yellow correspond to gene sets with a high number of genes shared with our list of DEG. The more purple the node, the less similarity between the gene set and our list of DEGs. The number of genes in each gene set is indicated inside each node. The thickness of the edges represents the similarity coefficient: the more similar two gene sets the thicker the edge that connects them. The nodes with a degree of 1 or 0 that were not connected to the network were removed. To define major biological clusters represented in these gene sets we used the tool AutoAnnotate which automatically defines clusters within the network. AutoAnnotate first clusters the network using the clusterMaker2 application in Cytoscape and draws a circle around each cluster. Then, an annotation associated with the three words more represented in the cluster is added using the WordCloud app. These annotations have been manually curated to ease data interpretation. The size of the cluster label reflects the number of nodes of that cluster.

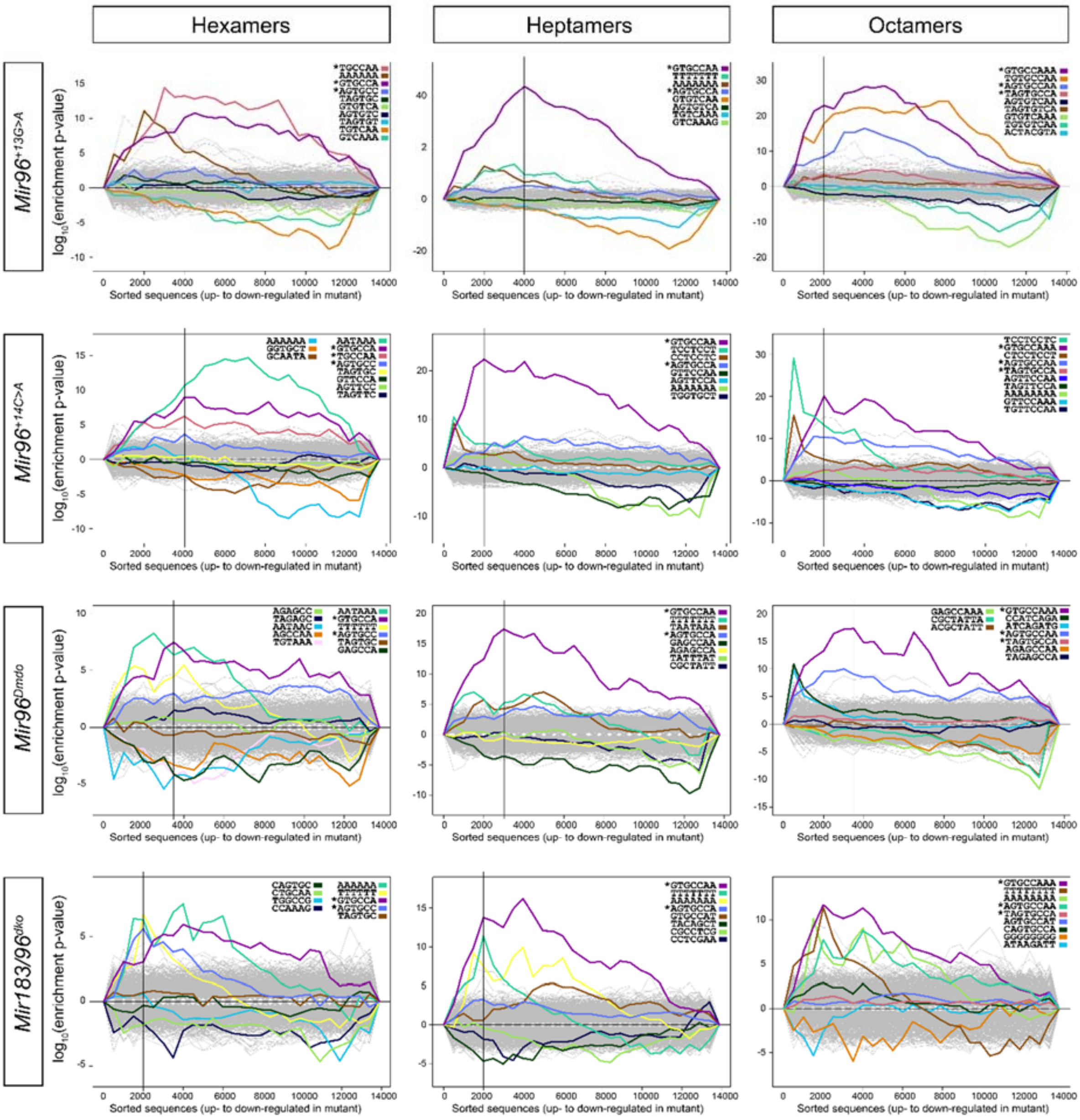


**Figure S2. Sylamer plots of hexamers, heptamers and octamers in the misregulatred transcriptomes of four *Mir96* mutants.** For *Mir183/96*$^{dko}$ and *Mir96*$^{Dmdo}$, the RNAseq data from (Lewis et al. 2020) and the microarray data from (Lewis et al. 2009) were used respectively, and replotted with 3'UTR data from GRCm39. Starred words indicate wildtype seed region matches. The vertical line on each plot shows the location of the peak enrichment of the wildtype seed(s), which was determined to be the first peak of a word corresponding to a wildtype seed region. Candidate targets are the genes to the left of this line (so the most upregulated genes) which have a wildtype seed region match in their 3'UTR.

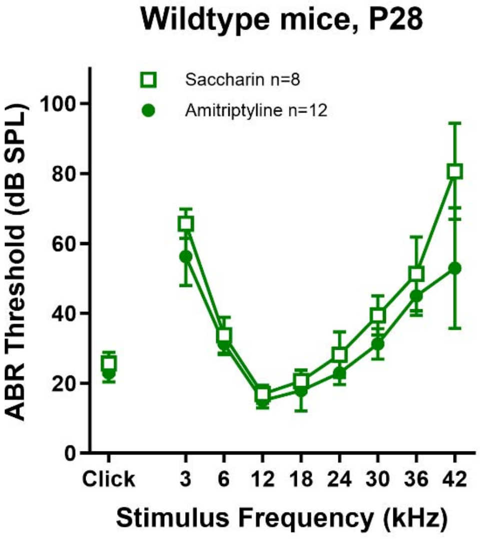


**Figure S3. Testing amitriptyline on C57BL/6N wildtype mice.** No difference in thresholds was observed between wildtype mice given 200µg/ml amitriptyline and 2% saccharin in their water and wildtype mice given just 2% saccharin at 4 weeks old.
